# Is there a pathological switch that triggers the onset of renal calcification?

**DOI:** 10.1101/2025.01.09.630316

**Authors:** Thamarasee M. Jeewandara

## Abstract

**Introduction:** Nephrocalcinosis, nephrolithiasis and Randall’s plaque formation are distinct renal pathologies of biomineralization predominantly originating in the renal papillae. Experimental evidence on the events leading to the initial aggregation of nanometer-scale plaque or stone deposits in these regions are limited. Cellular plasticity is a regulatory mechanism of disease progression, and can lead to the transition of epithelial to mesenchymal stem-cell-like phenotypes, and generate macrophages to trigger pathophysiological alterations underlying renal biomineralization. We aim to understand the pathological mechanisms of biomineralization at the renal papillary tip of clinical patient samples and develop functional assays to analyze mechanisms of disease progression within organ-chip devices *in vitro*.

**Methods:** We analyzed clinical cohorts of patient renal papillae tissues obtained via nephrectomy (n=34) categorized as stone formers (SF) vs. non-stone formers (NSF). We studied the histopathology and genetic (bulk RNA-sequencing) composition of patient samples in the two groups. We examined the role of primary cells, including peripheral blood mononuclear cells (PBMCs) - progenitors of macrophages, isolated from patient blood samples to differentiate M1 pro and M2 anti-inflammatory macrophage phenotypes for static culture and flow/stretch analyses on organ-on-a-chip devices (Emulate Inc). We stained tissue sections with histology dyes and conducted digital pathology multiplexing analyses via quantitative pathology software (quPath, GitHub) by training an artificial neural network. We conducted fluorescence in situ hybridization (FISH) studies to identify genetic biomarkers of inflammation extracted from the bulk-RNA sequencing data.

**Results:** Based on the initial results of digital pathology, we identified renal calcium deposits (p value = 0.0017), collagen deposits (p value = 0.0001), fibrosis (p value = 0.0385) and renal casts or inflammatory cells among SF vs NSF cohorts across the cortex-to-tip region of renal papillae. Bulk RNA-sequencing analyses were primarily conducted with DAVID-KEGG and Panther 17.0 classification databases to highlight key regulatory pathways of interest involved at the onset of renal biomineralization, such as the oxidative stress pathway, hypoxia response via HIF activation, and inflammation mediated by chemokine and cytokine signaling. The FISH studies identified genes involved with inflammation; GALNT3, PLEKHO1, SLCO2A1, and VCAM1. We successfully differentiated patient-derived PBMCs to M1 and M2 macrophage lineages to study the impact of oxidative stress by using static 35 mm plate and flow microfluidic organ-chip instruments, to conduct appropriate functional assays in cell culture.

**Conclusion:** The study outcomes provide insights to the precursors of renal biomineralization and delineated the expression of a pathological switch at the onset of hypoxia. The data will provide a fundamental framework to isolate primary cells from patient samples to conduct cell culture studies under static conditions, and translate the outcomes to flow analyses on a Kidney Chip instrument (Emulate. Inc) to mimic pathological conditions in a microphysiological environment *in vitro*. The ultimate outcome of this project will lead to the development of functional assays that emulate the kidney microphysiology on an organ-chip instrument, suited for clinical translation as a personalized, precision diagnostics and therapeutics platform.

## Introduction

In the spectrum of kidney ailments, chronic kidney disease (CKD) affects the structure and function of kidney filtration including nephron tubule portions to cause the build-up of kidney filter wastes and excess fluids from blood, leading to dangerous levels of fluid, electrolyte and waste buildup in the body (<u>Levey A. et al. 2012</u>). Renal tissue mineralization (nephrocalcinosis), stone formation (nephrolithiasis) and Randall’s plaque formation are distinct renal pathologies that contribute to stone formation by originating in diverse regions of the kidney, with limited information available on the events leading to the initial aggregation of nanometer-scale plaque or stone deposits (<u>Wiener S. et al. 2019</u>, <u>Ho S. et al. 2018</u>). Typically stones that form in the renal collecting system are attached to a calcium phosphate lesion of the renal papillae known as Randall’s plaque, while stones can also form in the duct of Bellini, or in free solution (<u>Coe F. et al. 2010</u>). How renal calcification originates at the tip of the renal papillae is a century-old puzzle.

The mechanism of human kidney stone formation can be classified into four categories - Growth over Randall’s plaque as seen in idiopathic calcium oxalate (CaOx) stone formers where most calculi are formed on Randall’s plaque. Growth across Bellini duct plugs seen mostly in calcium oxalate and calcium phosphate formers. And microlithiasis formation that is apparent in the inner medullary collecting ducts observed during cystinuria, and the formation in free solution within renal calyces or the renal collecting system, as observed in calcium oxalate stones in patients with hyperoxaluria, and brushite or hydroxyapatite stone formers that constitute the core of Randall’s plaque (Jeewandara T. et al. 2023, <u>Rao C. et al. 2019</u>, <u>Evan A. et al. 2014</u>). Oxidative stress affects the first two mechanisms via endothelial injury during progressive stone formation (<u>Jeewandara T et al. 2024</u>, <u>Saenz-Medina J et al. 2022</u>).

In the present study, we analyzed clinical cohorts of patient renal papillae tissues (n=34) obtained via nephrectomy, categorized as stone former (SF) vs. non-stone formers (NSF). We studied the histopathology and genetic bulk RNA-sequencing composition of patient samples in the two groups. The work addressed a key question ‘Is there a pathological switch at the renal papillary tip that triggers the onset of calcification?’ The puzzle of how cells sense and adapt to oxygen availability is an age-old question that eventually led to unveiling its vital biological phenomenon with pioneering research (<u>Nature Collection 2019</u>, <u>Forsythe J. et al. 1996</u>, <u>Schofield et al. 2004</u>). Preceding work showed that oxygen sensitive enzymes and cellular machinery can come together to “turn off” a major regulatory protagonist known as the hypoxia inducible factor 1α (HIF1α) proteins (<u>Chopra A et al. 2020</u>). In our work, we show how the very same protagonist (HIF1α) is at play during hypoxia-driven renal calcification; substantiated with -omics driven data, histopathology, and functional assays to support our hypothesis of the presence of a biological switch that can regulate calcification at the renal papillary tip region in this study. In the present study, we established a dynamic microphysiological system by using microfluidic devices replete with patient-derived and differentiated macrophage cell types to recreate a pathological microenvironment to conduct several bioinspired functional assays (Figure 1). In essence, we bottom-up engineered a hypoxia-driven pathological cell signaling pathway on a kidney-on-a-chip instrument based on first-principles of tensional integrity or tensegrity (<u>Ingber D. E. 2020</u>, <u>Ingber D.E. 1993</u>). Unless otherwise stated all experiments in this study were conducted with clinical samples obtained from both stone former vs. non-stone formers. We investigated our study outcomes across four main aims –

- Histopathology studies at the renal papillary tip to understand the pathological mechanisms of renal calcification *in vivo,*
- The transcriptomics and gene-specific outcomes at the renal papillary tip to explore mechanisms of renal calcification,
- Conducting functional assays that are informed via histopathology and transcriptomics analyses to recreate or emulate pathological mechanisms on a kidney-chip
- Using organ-on-a-chip platforms to recapitulate patient pathology, to develop state-of-the-art microphysiological platforms.

**Figure 1:**
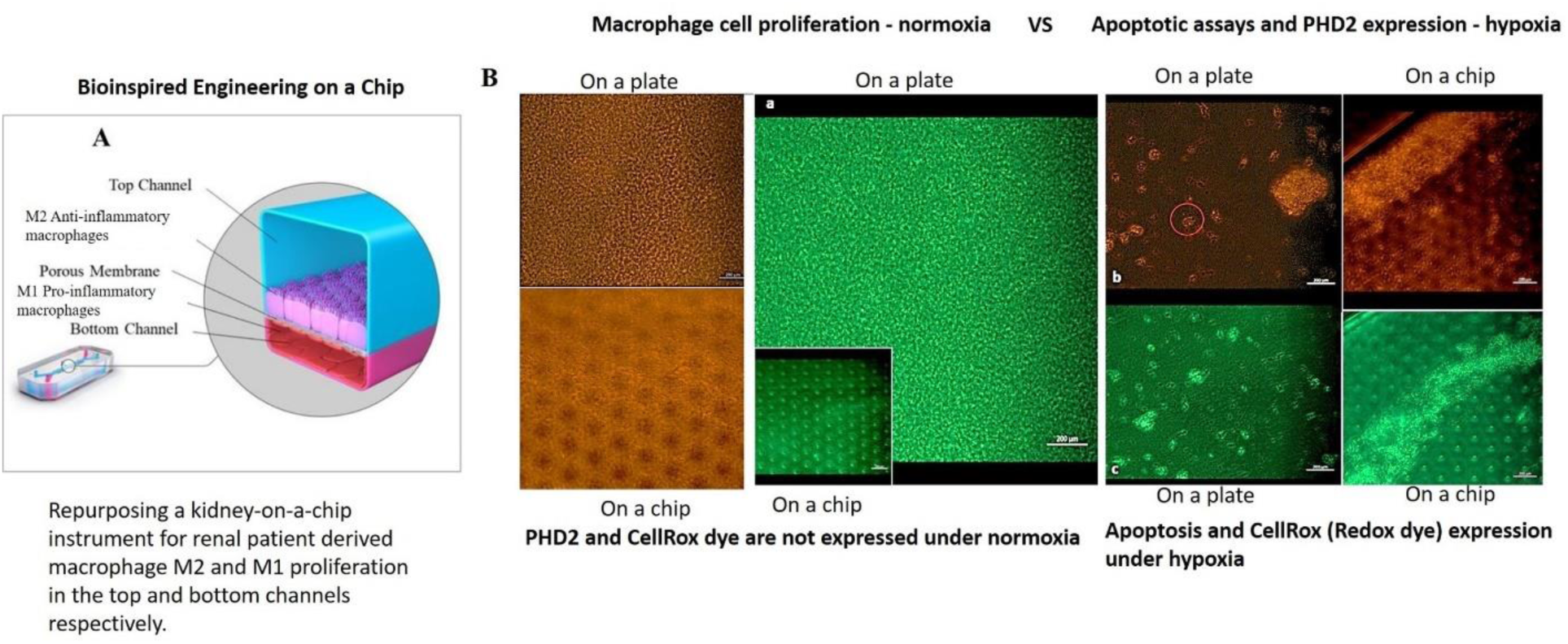
Study recap – recreating a pathological microenvironment on a kidney-on-a-chip microfluidic instrument to investigate the presence of a biological switch underlying renal calcification. A) Repurposing a proximal tubule-on-a-chip (kidney-on-a-chip) instrument for renal patient-derived macrophage M1 and M2 proliferation on the top and bottom channels of the organ-chip instrument. B) a-c: The confluence of the macrophage cell lines was well supported on the organ chip instrument surface and on the tissue-culture plates: the cluster-forming cells are motile. Apoptotic assays were conducted on a chip and a plate to emulate bioinspired oxidative stress to observe the expression of the biomarkers of prolyl hydroxylase domain 2 (PHD2) and CellRox dye (orange and green, respectively). The morphology of the cells dramatically changed from miniature, motile M1 and M2 variants to aggregated cell clumps emitting PHD2 and CellRox under hypoxia [Jeewandara T et al. 2023].

## Materials and Methods

### • Human specimen

The analysis cohort comprised of cross-sections of the renal papillary tip region of renal stone forming and non-stone forming patients, to represent n=34 total tissue samples obtained via nephrectomy at the UCSF Medical Center under protocols approved by the University of California San Francisco’s Committee on Human Research Protection Program (IRB no. 14-14533 and protocol no. H8933071801). All studies involving human specimen were conducted in accordance with the 1964 Helsinki declarations or comparable ethical standards.

### • Tissue preparation for histology – renal tip extraction from Randall’s plaque patients

Dulbecco’s phosphate buffered saline (DPBS, Sigma-Aldrich) (5 mL) was prepared with calcium and magnesium, supplemented with 1% penicillin-streptomycin in a 15 mL sterile conical tube to carry out tissue preparation for renal tip extraction (<u>Arbra C. et al. 2018</u>). Ice was placed next to the operating table and the kidney tissue was placed in 1% penicillin-streptomycin within the 15 mL of DPBS on ice. The kidney section was transferred from a 15 mL conical tube to a 60 mm petridish and rinsed twice using 5 mL DPBS with calcium and magnesium, and maintained in 5 mL DPBS in a 60 mm dish (<u>Arbra C. et al. 2018</u>). In an additional 60 mm petridish, 5 mL of DPBS was added for the microdissection protocol. The 60 mm petridish with the kidney section in DPBS was incorporated from the tissue culture hood to the microscope imaging area. At a magnification of 3.2 x, we dissected an inverted pyramid tissue from the area of interest at a volume of 2 mm^2^ in size. After decreasing the magnification (1.5 – 2.0 x) for the remainder of the section, the tissue slice was fragmented into progressively smaller pieces, until the tissue segments were less than or equal to the diameter of the needle tips. A tissue volume of 2 mm^2^ produced more than 50 segments upon complete dissection. After dissection, the 60 mm dish with renal segments were moved to the tissue culture hood to remove DPBS with a 1000 µL pipette. These tissue samples were then fixed in 4% paraformaldehyde and processed for histopathology via paraffin embedding and tissue sectioning.

### • Histology staining and digital pathology – Training an artificial neural network (ANN) for quantitative pathology of human specimens

The tissue sections of the medullopapillary complex were fixed and paraffin-embedded for histopathology staining protocols (Table 1), at the Zuckerberg San Francisco General Hospital. An artificial neural network was trained at the Biological Imaging Development CoLab, UCSF by using a quantitative pathology software (quPath, GitHub) to quantify the presence of key biomarkers of renal calcification, including tissue areas of vimentin deposits, calcium deposits, and collagen sedimentation within the renal tissues of stone former vs. non-stone former cohorts. During the image feature extraction process, to train the software, we used n=17 patient samples, where SF = 9 samples and NSF = 8 samples each, pertaining to each dye of interest (Table 1). Regions or morphological features of interest were identified for a given stack or collection of biomedical images stained with a histopathology dye of interest, and each tissue region/zone were classified using the quPath software by loading a pixel classifier to extract information from the image. The pixel classifier was further trained on script editor of the software to identify regions of the histology image collections (including vimentin, calcium, and collagen dyes). The specific areas of interest were quantified by choosing the ‘artificial neural networks’ classification model for image segmentation, to identify tissue regions from the cortex-to-the-tip of the renal papillae and labelled Z1-to-Z4 (Jeewandara T. et al. 2023, <u>Bankhead P. et al. 2017</u>, <u>Ho S. et al. 2018</u>). To optimize the quantification, the ‘create objects’ option of the software was selected, and specific morphological regions of interest relevant to a minimum object size of the image were defined. A thresholder was created to analyze each image, the steps for data collection were automated by writing up a macro. The extracted data was transferred via Excel for multivariate analysis.

**Table 1:** A summary of histopathology stains used to identify and quantify regions of interest in renal papillary tips of stone formers vs. non-stone formers by training an artificial neural network.

| Pathology stain | Region of interest identification |
| --- | --- |
| Alizarin Red | Renal calculi or renal calcification seen as red deposits |
| Vimentin-DAB | Epithelial-mesenchymal transitions seen as brown deposits |
| Hematoxylin and Eosin | Identification of inflammatory deposits |
| Masson’s Trichrome | Collagen/muscle identified as green and purple regions |

### • Histopathology with renal tissue – multivariate analysis

The tissue regions of interest were quantified relative to the uptake of different histopathology dyes (Table 1) using the ANN-derived data that were extracted for multivariate analysis. The log values of all tissue sample regions of interest were obtained and quantified via ANN pertaining to 1) tissue areas of calcium deposition (stained red), 2) tissue areas of epithelial-mesenchymal transitions (stained brown), and 3) tissue areas of collagen deposition (stained green) across all renal papillary cross-sections – from the cortex to the tip region (Z4 to Z1). Statistical analysis of the data (GraphPad Prism) determined the *p*-values. The statistical significance of the biomarkers of interest were derived in the renal medullopapillary tissue regions between the stone former vs. non-stone former patient cohorts. Differences with a *p-* value < 0.05 were considered significant.

### • Transcriptomics with renal tissue – bulk RNA sequencing and enrichment assays

For the RNA sequencing study, papillae were snap frozen in liquid nitrogen shortly after collection, and stored at -80^0^C until further use. Frozen samples were placed on a petridish on ice and allowed to soften, the papillae were sectioned longitudinally from the tip to the outer medulla to prepare thin slices of the whole papillae that were then transferred into a Dounce homogenizer. The lysate was transferred into a shredder spin column and centrifuged at 16,000 g for 2 minutes. The supernatant was extracted to purify RNA using the RNeasy Plus Universal Mini kit (Qiagen) according to the Manufacturer’s protocol. The RNA concentration was quantified using a Nanodrop spectrophotometer and Qubit fluorometer to identify DNA contamination. The quality control check of the RNA was performed by running 2 µL of 10 ng RNA on a fragment analyzer according to the manufacturer’s protocol. Libraries were prepared from 300 ng of RNA with universal plus mRNA-sequencing library preparation kit (Tecan). The pool of libraries were loaded onto MiniSeq (illumina) according to the protocol. The normalized libraries were sequenced on one lane of HiSeq 4000 SE50 to yield 35 M reads per specimen. Reads uniquely mapped to known mRNAs were used to assess expression changes between genes.

### • Bioinformatics – KEGG pathway analysis and GO term enrichment analysis

Pathway enrichment analysis was performed using the Gene ontology (GO) and Kyoto Encyclopedia of Genes (KEGG) database resources. The analyses revealed functional gene sets that significantly clustered under 64 pathological pathways. The read counts of the upregulated genes from bulk RNA-sequencing data were normalized and used to run KEGG pathway analysis and Gene ontology enrichment analyses. Both analyses were performed using the Database for Annotation Visualization, and Integrated Discovery (DAVID) (<u>Ashburner M. et al. 2000</u>) and Panther 19.0 classification system with large-scale functional annotation gene lists from genome-wide data that were gathered from bulk-RNA sequencing experiments, from molecular to cellular to the organism-level systems (<u>Mi H et al. 2013</u>, <u>Mi H. et al. 2019</u>). The input data consisted of gene/compound information and specific target pathways and the outcomes generated a native Kyoto Encyclopedia of Genes and Genomes view to represent all meta-data on pathways, including spatial and temporal information, tissue/cell types, inputs, outputs, and connections. A comprehensive heatmap was generated from the bioinformatics data obtained from kidney patient renal papillary tip tissues to segregate the genes expressed between stone formers and non-stone formers. Using Library Pathview, an R-package for pathway-based data integration and visualization, we obtained maps of user data (selected pathways that were upregulated/downregulated) as relevant pathway graphs (<u>Yu G. et al. 2012</u>). The dot-plots showed the activity of enriched signaling pathways to view genes in the context of biochemical cascades. The enrichment dot plots were developed using an R-package to compare biological themes among gene clusters to interpret the omics data.

### • Lymphoprep – peripheral blood monocyte cell separation

Blood samples were obtained from stone formers vs. non-stone formers in collection tubes (n = 10). The blood from the separate vials were added into a 50 mL flask and PBS added to dilute to 20 mL in total. Lymphoprep (15 mL, StemCell Technologies) was carefully added to the bottom of a new 50 mL conical flask. Slowly, 20 mL of the diluted blood was pipetted on top of the Lymphoprep. The tubes were spun at 800 rpm for 25 minutes at room temperature with a slow start for 30 minutes. The top layer of media (yellowish, containing the PBMCs) was aspirated without disturbing the mononuclear layer. The mononuclear layer was harvested (cloudy white) between the media and Ficoll by using a 10 mL pipet and placed into a new 50 mL container. The collection media was minimized above, with the Lymphoprep below, without drawing red blood cells from the erythrocyte pellet. A new conical flask was filled with 30 mL of PBS and spun at 800g for 10 minutes. The media was gently aspirated without disturbing the pellet, until 3 mL remained. A volume of 1-2 mL RBC lysis buffer was added and swirled gently for 1-2 minutes. The container was filled to 15 mL to spin again at 800 g rpm for 10 minutes, the media was aspirated without disturbing the pellet and the pellet resuspended in freezing buffer (Bambanker, 1 mL) to immediately freeze or in working buffer.

### • Cell culture – M1 and M2 Macrophage differentiation

The differentiated blood mononuclear cells/monocytes were isolated and plated in monocyte attachment media (MAM, PromoCell) in static 35 mm cell culture plates. After 1.5 hours, the medium was aspirated along with floating non-adherent cells. The adherent cells were washed 3x with monocyte attachment medium. The cells were immediately incubated with the complete M1 and M2 macrophage generation medium (PromoCell), and left to grow for 6 days. By day 6, another 50-75 percent by volume of fresh completed M1 or M2 macrophage generation medium DXF (PromoCell) was added. After 1-day, appropriate activation or polarization factors were added as cytokines to facilitate M1 pro-inflammatory (TNF-α, IL-17 A) and M2 anti-inflammatory (IL-4, IL-13) macrophage phenotypes (PromoCell). By day 2, the medium was changed to aspirate and collect the floating cells. Further complete M1 and M2 macrophage generation medium DXF was added to recover floating cells and combined with the culture. By day 10, the macrophages were ready to use, or could remain in culture. After day 10, the macrophages were either harvested or sub-cultured by using macrophage detachment solution DXF (PromoCell) and the cells were prepared for quality control analysis using flow cytometry.

### • Cell culture – Harvesting and sub-culturing macrophages

The M1 and M2 macrophages that were differentiated from mononuclear peripheral blood cells were validated or identified via cell-sorting with fluorescence-activated cell sorting (FACS). The culture medium of M1 and M2 macrophages was aspirated on day 10, and the cells were washed twice with PBS. Immediately, an appropriate amount of cold macrophage detachment solution (DXF) (PromoCell) was added (e.g. 25 mL per T-75 flask) and the sealed tissue culture vessel was incubated for 40 minutes at 2-8^0^C. The cells can be incubated for another 20 minutes, if necessary, to enforce cell release from the tissue culture plastic and the organ-on-a-chip instruments. Cells being prepared for FACS were not scraped to dislodge the remaining macrophages from the tissue culture plastic. The harvested macrophages were collected in centrifugation tubes and diluted with PBS (1:1) or ice-cold fluorescence-activated cell sorting buffer (PBS 0.5-1% BSA or 10% FBS) and centrifuged for 15 minutes at 350 g, at room temperature, followed by cell counting. The macrophages (105 per sample at 100 µL) were collected in four, 1 mL centrifugation tubes for immunofluorescence assays for fluorescence-activated cell sorting preparation.

### • Quality assurance – flow cytometry with fluorescence-activated cell sorting (FACS)

A blocking antibody-step was conducted to prevent high level of cellular expression of Fc receptors that may contribute to non-specific binding and background fluorescence. Blocking buffer (100 µL, Cell Signaling Technology) was added into each sample (1:50 ratio) and incubated on ice for 20 minutes and centrifuged at 1500 rpm for 5 minutes at 4^0^C. The supernatant was discarded. The M2 cells were incubated with mouse monoclonal CD163 antibody (1:100, ThermoFisher Scientific) and stored at 0.5 hours at 4^0^C. The M2 cells were again incubated with a light-sensitive, fluorescent goat anti-mouse polyclonal IgG antibody (AlexaFluor 647, abcam) at a dilution of 1:200 for 0.5 hours at 4^0^C or left on ice. The cells were washed by centrifugation at 1500 rpm for 5 minutes and resuspended in 200 µL-to-1 mL of ice-cold FACS buffer, and stored on ice or at 4^0^C in a fridge, until the scheduled time for analysis to prepare the cells for flow cytometry. For M1 cells, the steps were carried out similarly, and the cells were incubated with mouse monoclonal CD86 (1:100, ThermoFisher Scientific) and stored for 0.5 hours at 4^0^C on ice. The cells were washed by centrifugation at 1500 rpm for 5 minutes, and the supernatant was discarded. The cells were incubated with a light-sensitive, fluorescent goat anti-mouse polyclonal IgG antibody (AlexaFluor 647) at a dilution of 1:200 for 0.5 hours at 4^0^C, covered by foil on ice. The cells were washed by centrifugation at 1500 rpm for 5 minutes, resuspended in 200 µL-to-1 mL of ice-cold FACS buffer (approximately 500 µL) and stored in a fridge at 4^0^C or on ice until scheduled time for analysis.

### • Repurposing a proximal-tubule-on-a-chip to study renal biomineralization

The kidney-chip compartment that emulates the proximal kidney tubule on a chip were seeded with macrophage M1 and M2 cells derived from peripheral blood mononuclear cells of renal stone forming vs. non-stone forming patients, to differentiate and culture cells in lab. The instrument was repurposed to mimic the renal papillary tip region. The cells were cultured on cell-specific extracellular matrix proteins, and maintained in static culture for up to four days prior to connecting the culture chip to the Zoë instrument, to facilitate continuous cell culture media flow. Cell culture experiments were conducted in accordance to the Emulate protocol (Emulate EP177 V1.0) by initially preparing a proprietary ER-1^TM^ surface activation reagent (5 mg) under UV light for five minutes. The solution was introduced to the channels to activate them for fifteen minutes, and the chips were washed. The extracellular matrix solution was prepared with Matrigel (100 µg/mL) by thawing the solution overnight at 2-to-6-degree Celsius and with collagen IV (30 µg/mL) by dissolving the powder in PBS (5 mL). The prepared extracellular matrix solution was aliquoted to a desired volume (100-200 µL) for the coating step of the organ-chip instrument. After completing the macrophage differentiation and cell detachment steps above, the cells (100-200 µL) were introduced into the organ-on-a-chip instrument and to the 35 mm cell culture dishes for their growth. The macrophage M1 cells were plated in the bottom channel at a density of 2 x 10^6^ cells/mL and the chips were inverted for 3 hours to facilitate attachment to the porous membrane that separates the two channels in the chip. The top channel was seeded with M2 macrophage cells at a density of 1 x 10^6^ cells/mL. A gravity wash was conducted with fresh media after cell attachment to the instruments to ensure nutrient replenishment, approximately 3 hours post-seeding. The cells were washed the next day and for the period of culture with the culture medium, and connected to the pods and to the Human Emulation System (Emulate Inc). The flow rate was set at 60 µL/hour for the top and bottom channels and the medium replenished appropriately for the duration of the experiment.

### • Functional assays – oxidative-stress initiated apoptosis – Redox reactions

M1 and M2 cell cultures were subject to oxidative-stress initiated apoptosis by inducing chemical ischemia with sodium dithionate in liquid nitrogen for detection with CellRox redox dye (Life Technologies). A hypoxic buffer was prepared with 4 mM final concentration of sodium dithionite in oxygen glucose deprived (OGD) physiological buffer. The media was exposed to liquid nitrogen for 10 minutes to achieve OGD, prior to dithionite addition. If frozen upon exposure to liquid nitrogen, the OGD solution was thawed and warmed up to room temperature. Sodium dithionite (4 mM, Sigma-Aldrich) was added to the OGD in each 35 mm cell culture dish (1 mL) or microfluidic chip (200 µL) holding the cells to induce rapid and reliable hypoxia-like conditions, and incubated for 30 minutes. A protein blocking buffer (1000 µL per well, and 200 µL per chip, abcam) was added for 30 minutes-to-1-hour to block non-specific protein binding. The CellRox dye reagent (2.5 mM) was added to the 35 mm culture dish (10 µL) or kidney-chip (1-5 µL). The cells were incubated for 30 minutes-to-1-hour. The cells were washed twice with PBS and incubated with DAPI (1:1000) at 100 µL per plate or chip. The cells were washed with PBS twice again, fixed with 10% neutral buffered formaldehyde, washed with PBS, and observed under live microscopy. Images were captured using Zeiss (Zeiss Observer Z1 Axiocam 506) in AG520 channels to capture images expressing the protein of interest.

### • Functional assays –prolyl hydroxylase domain 2 (PHD2) expression

Two separate cell cohorts (of M1 and M2 cell cultures) were used to analyze the positive vs. negative expression of PHD2 via apoptotic functional assays (n=10). The cells were subjected to chemically induced ischemia by preparing a hypoxic buffer with a concentration of sodium dithionite (4 mM, SigmaAldrich) in oxygen glucose deprived (OGD) physiological buffer. The media was exposed to liquid nitrogen for 10 minutes to achieve OGD, prior to dithionite addition. If frozen upon exposure to liquid nitrogen, the OGD solution was thawed and warmed up to room temperature. Sodium dithionite (4 mM) was added to the OGD in each 35 mm cell culture dish (1 mL) or microfluidic chip (200 µL) holding the cells to induce rapid and reliable hypoxia-like conditions, and incubated for 30 minutes. A protein blocking buffer (1000 µL per well, and 200 µL per chip) was added for 30 minutes-to-1-hour to block non-specific protein binding. After removing the buffer, PBS was added with the primary antibody (EGLN1/PHD2 1:50 dilution, Novus Biologicals) for incubation; prepared at 50 µL per well for three wells. The samples were incubated for 1 hour in a warm incubator and washed with PBS. The secondary antibody, anti-rabbit raised in goat (1:200) (Goat primary antibody to Rabbit IgG-Alexa Fluor 555, abcam) was added to the wells/chips and incubated for 1 hour in a warm incubator and washed with PBS thereafter. The cells were washed twice with PBS and incubated with DAPI (1:1000, ThermoFisher Scientific) at 100 µL per plate or chip. The cells were washed with PBS twice, fixed with 10% neutral buffered formaldehyde, washed with PBS again and observed under live microscopy. Images were captured using Zeiss (Zeiss Observer Z1 Axiocam 506) in AF555 channels to capture images expressing the protein of interest.

### • Fluorescence in situ hybridization (FISH)

Stone former and non-stone former papillae were fixed in 10% neutral-buffered formalin for 18-24 hours. Upon fixation, tissues were immediately processed and embedded in paraffin. FISH was performed with RNAscope multiplex fluorescent detection kit V2 according to the manufacturer’s protocol. Kidney papillary tissue (4 µm) sections were hybridized with target probes for specific genes of interest (Table 2) and labelled with Opal fluorophores (Opal 520, 570, 620, 690). The nuclei were counterstained with DAPI and sections were mounted using ProLong Gold Antifade (Invitrogen). Images were captured using Zeiss (Zeiss Observer Z1, Axiocam 506) in FITC, CY5, TexasRed and Cy5.5 channels to capture images of the papillary tissue and evaluate the differences between the gene expression levels in the tissue regions by using MATLAB. The Otsu algorithm (<u>Otsu N. 1979</u>) was applied to each grayscale image to determine the intensity threshold. Differences with a *p-*value < 0.05 were considered significant.

**Table 2:** The kidney papillary tissue sections were hybridized with target probes for specific genes of interest obtained from Advanced Cell Diagnostics.

| Gene | Gene probe | Cat number | GenBank Accession |
| --- | --- | --- | --- |
| SLCO2A1 | RNAscope™ Probe HS VCAM1 | 467691 | NM_005630.2 |
| LOX | RNAscope™ Probe- Hs-LOX | 415941 | NM_001178102.1 |
| PLEKHO1 | RNAscope™ Probe- Hs-PLEKHO1-C2 | 874501-C2 | NM_001304723.1 |
| VCAM1 | RNAscope™ Probe- Hs-VCAM1-C2 | 440371-C2 | NM_001078.3 |
| IGF1 | RNAscope™ Probe- Hs-IGF1-C3 | 313031--C3 | NM_000618 |
| TMEM30B | RNAscope™ Probe- Hs-TMEM30B-C3 | 874471-C3 | NM_001017970.3 |
| GALNT3 | RNAscope™ Probe- Hs-GALNT3-C4 | 874481-C4 | NM_004482.4 |
| HAPLN3 | RNAscope™ Probe- Hs-HAPLN3-C4 | 874511-C4 | NM_001307952.2 |

### • Statistics

The image processing data were extracted from the quPath image quantification software (quPath, GitHub) as mean values to Office Excel and graphed in GraphPad software (San Diego, California, USA), and their *p* values were quantified and represented. Differences with a *p-*value < 0.05 were significant for all study components. The histopathology imaging data were analyzed using multivariate analyses with Office Excel, and two-way ANOVAS were conducted with multiple comparisons for >2 variables using GraphPad software (San Diego, California, USA) made available for Windows PC.

## Results

### • Digital pathology – training an ANN and conducting multivariate analysis of pathology data

The initial outcomes of the artificial neural network (ANN) assisted quantitative pathology (quPath, GitHub) measurements were quantified for stone former vs. non-stone former samples that were obtained via nephrectomy from patient cohorts. The multivariate analyses indicated the presence of renal calcium deposits (p = 0.0017), collagen deposits (p = 0.0001), and renal fibrosis quantified with vimentin (p = 0.0385) (Figure 2). We additionally noted the presence of renal casts or inflammatory cells from across the cortex to the tip region of the renal papillae (Z1-to-Z4). As expected, the outcomes of calcium deposition, collagen deposition due to fibrosis, and vimentin deposition due to epithelial mesenchymal transition indicated a significantly higher aggregation of the biomarkers in stone forming patients, when compared to non-stone formers (Figure 3). In general, the area of collagen was significantly higher when compared to the area of calcium and to the area of vimentin, respectively. To train the ANN, we used a classification strategy in the lab for the first time, to identify specific morphological features of the renal papillary tip, from the cortex-to-the tip region and developed a macro to automate the work process. We classified a cross-section of the renal papillae into four zones ranging from the cortex to the renal tip and identified the key morphological regions of the four zones to standardize the classification. We input morphological data to train the ANN for automated image analysis and facilitated high fidelity physical models to extract information and quantify regions from the image data.

**Figure 2:**
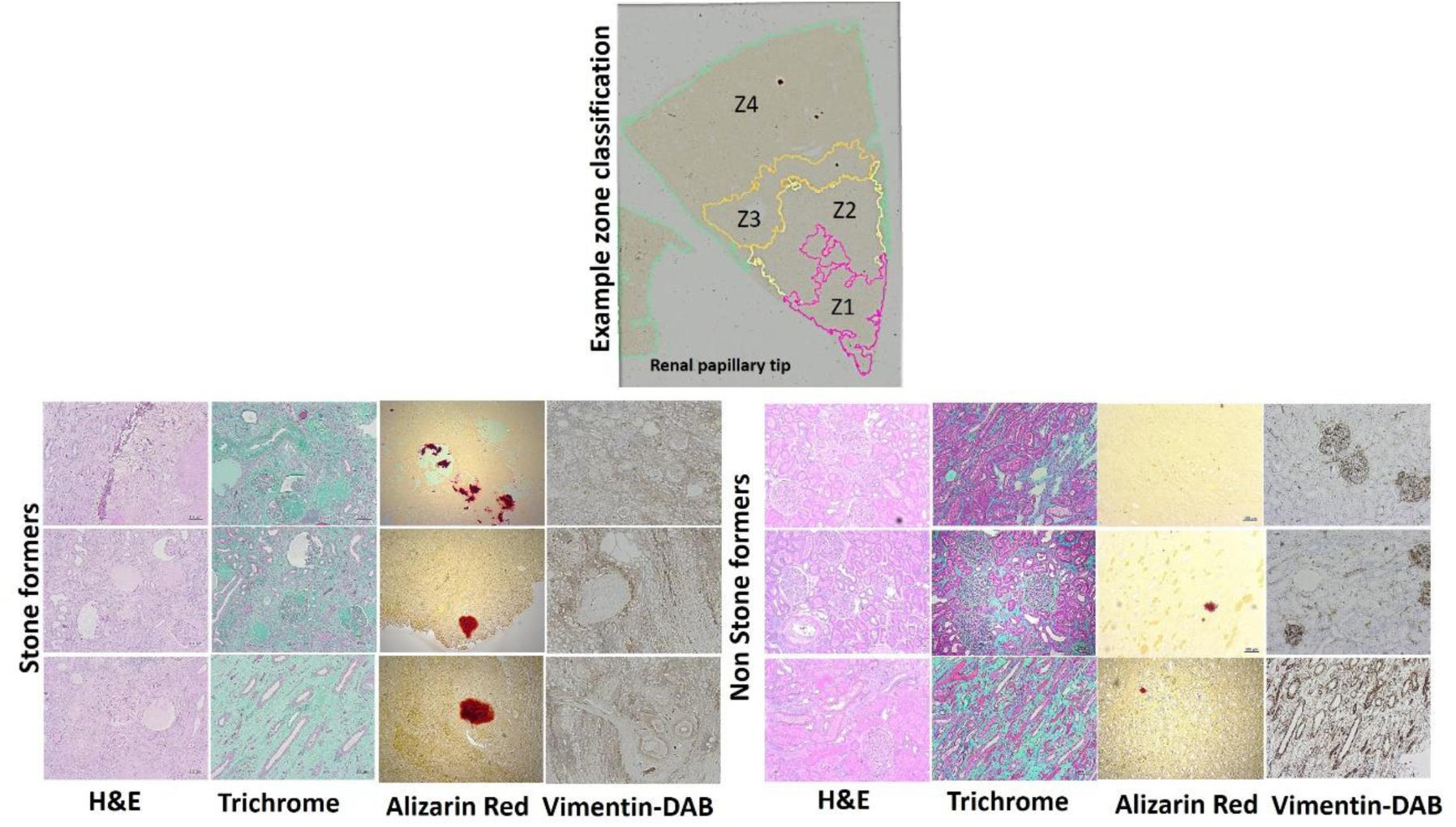
Histopathology data with stone formers vs. non-stone formers, the top image shows the method of zone classification that we used to train an artificial neural network with quPath software to quantify regions of interest (Z1-Z4) in the renal papillae. The tissue sections were stained with H&E to view inflammatory biomarkers in stone formers, including renal casts and epithelial thinning with a ropey or beaded appearance. In renal papillary sections of stone formers, the trichrome dyes indicated a higher percentage of green for increased collagen deposits, Alizarin red stained a higher percentage of crimson for calcium deposits, and vimentin DAB stained a higher percentage of brown to represent epithelial mesenchymal transitions during renal nephrolithiasis and showed the presence of glomerular amyloid balls in stone formers [Jeewandara 2023].

**Figure 3:**
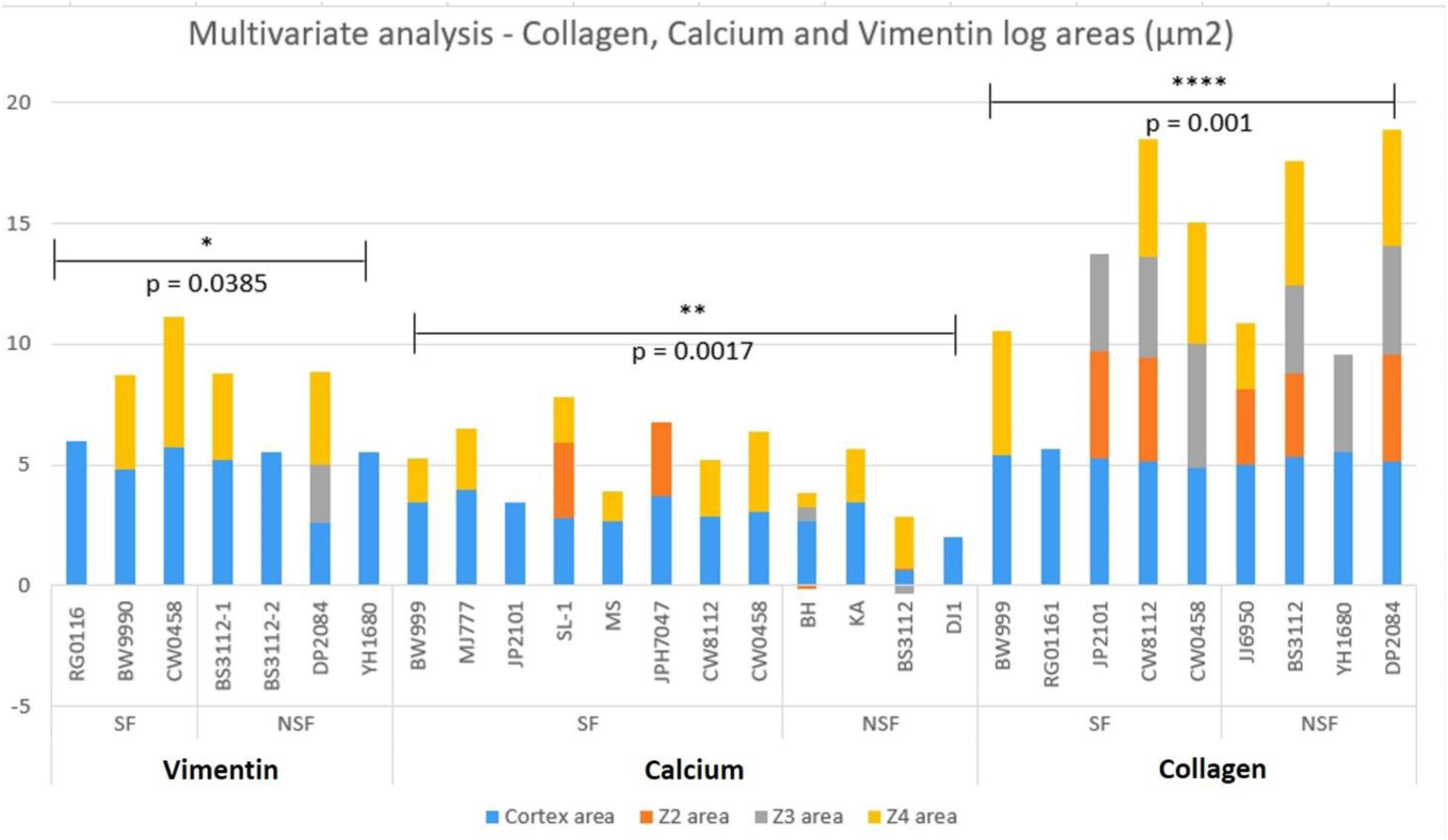
The quantified log values of the multivariate analyses of all samples; includes epithelial – mesenchymal transitions identified with vimentin DAB, calcium identified with alizarin red, collagen identified with trichrome, across the renal papillary cross-sections, and their p values. In this graph of ANN-derived classifications, the cortex area is assigned blue, Z2 area is orange, Z3 is gray, and Z4 is yellow. Not all tissue sections contained all classifications. There was significant variation observed in the data between stone formers vs. non-stone formers. Non-stone formers representing renal carcinoma showed higher variability with upregulated biomarkers of renal fibrosis (collagen) [Jeewandara T. et al. 2023].

**Figure 4:**
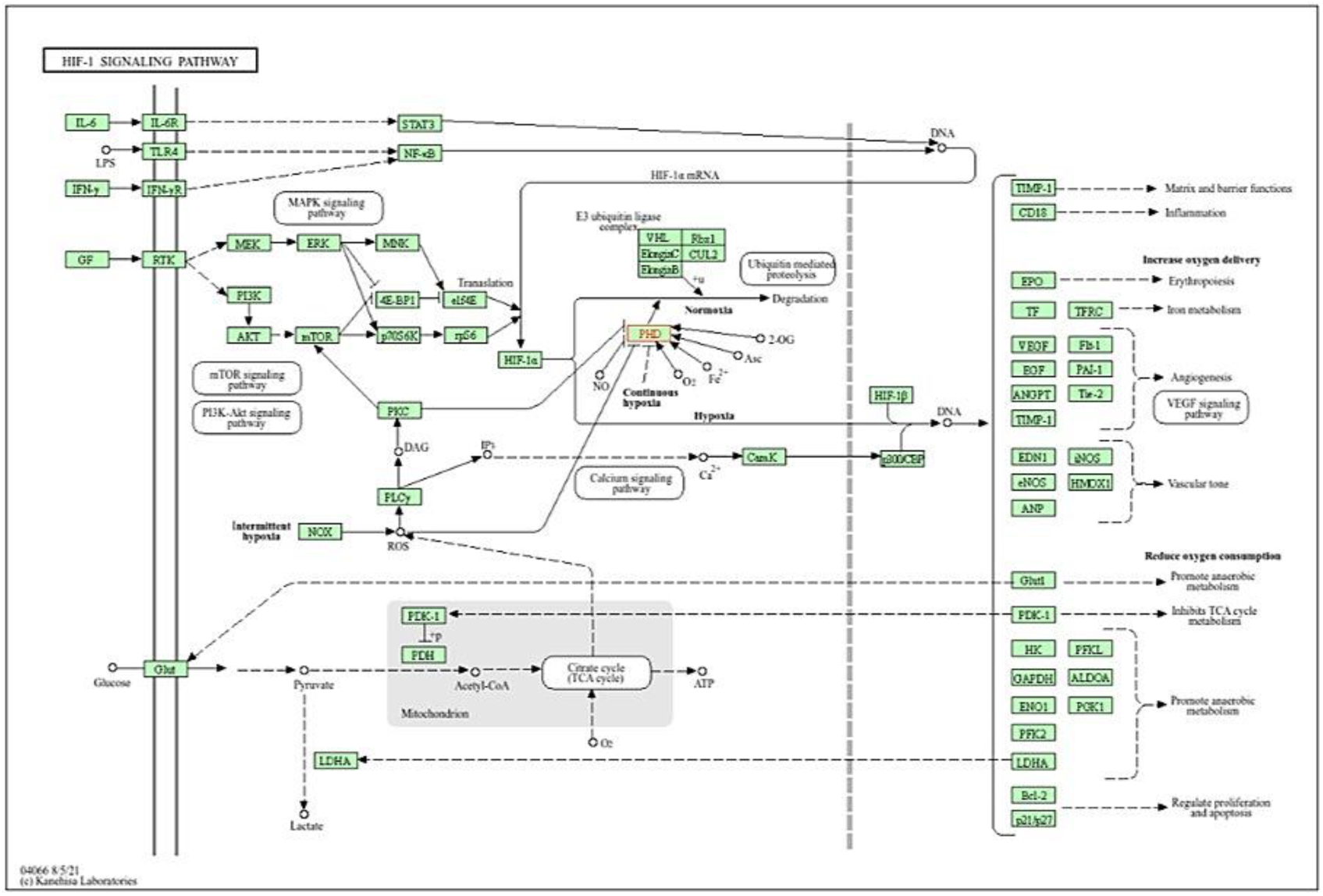
The KEGG meta-data pathway generated from patient-derived transcriptome bioinformatics data to elucidate key cascades of renal calcification – the image depicts a key pathway of interest in this study, the HIF1α signaling pathway associated with hypoxia-driven cell apoptosis during nephrolithiasis [<u>KEGG database</u>].

### • Transcriptomics and Bioinformatics - bulk RNA sequencing and enrichment dot- plots of the genes expressed at the renal papillary tip

For bulk RNA sequencing, we generated a heatmap and compared the gene expression of stone former vs. non-stone former patient papillary samples, to examine specific genes that are expressed and upregulated between the two patient cohorts [<u>Nookala A. et al. 2022</u>]. Differential gene expression (DE) was performed between the two patient cohorts. Of the 58,040 genes annotated in the genome, 1072 genes were significantly differentially expressed. Of these 765 were upregulated and 307 were downregulated. We used databases to visualize the key regulatory pathways associated with the genes expressed during renal calcification, including GO/DAVID-KEGG/Panther, as well as the Jackson Laboratory databases. The input data for the gene ontology (GO) database consisted of gene/compound information and specific target pathways, and the outcomes generated a native Kyoto Encyclopedia of Genes and Genomes (KEGG) view representing all meta-data on pathways, including spatial and temporal information, tissue/cell types, inputs, outputs, and connections. The DAVID-KEGG pathways highlighted key pathological cascades that contribute to renal calcification. We identified a total of 64 regulatory pathways of which the following 10 paths were highlighted in the study (Table 3). We selected a single pathway of ‘hypoxia response via hypoxia-inducible factor HIF1α activation,’ to conduct subsequent functional assays by bottom-up engineering the pathological cell signaling pathway on a kidney-on-a-chip instrument *in vitro,* to understand their influence on the trajectory of renal calcification *in vivo*.

**Table 3:** The pathological cascades of interest identified via database analyses of bioinformatics, and their mechanisms of interest recognized for further investigations to bottom-up engineer a pathological cascade on a kidney-on-a-chip microphysiological environment [Jeewandara T. et al. 2023].

| Pathological cascade of interest | Mechanism of interest during kidney stone formation | Reference |
| --- | --- | --- |
| Oxidative stress pathway | A primary mechanism underlying renal calcification | Saenz-Medina J et al. 2022 |
| Hypoxia response via HIF activation (core theme of the study) | Interactions of hypoxia-driven HIF1α and the switch-like activation of PHD2 that underly the onset of pathological calcification | Arsenault P. et al. 2016 |
| Transforming growth factor $\beta$ signaling pathways | Associated with cytokine upregulation in kidney tissue undergoing fibrosis and renal injury at renal biomineralization. | Khan S. et al. 2013 |
| Phagosomes | Associated with professional phagosomes such as macrophages | Russel D. et al. 2010 |
| FcγR mediated phagocytosis | Immune receptors that are primarily located on macrophages to clear calcium deposits during renal calcification | Vidarsson G. et al. 2014 |
| Natural killer cell mediated cytotoxicity | Associated with immune responses for M1 pro-inflammatory macrophage activation | Turner J. et al. 2019 |
| Actin cytoskeleton regulation. | Associated with cellular and molecular mechanisms during kidney fibrosis | Duffield J. et al. 2014 |
| Hedgehog signaling | Associated with cell proliferation and stem cell maintenance | Zhou D. et al. 2017 |
| P38 MAPK (mitogen-activated protein kinase) | Associated with cell stress and activating a macrophage-mediated inflammatory response | Drobnik M. et al. 2024 |
| JAK-STAT (Janus Kinase Signal Transducer and Activator of Transcription) | Associated with cell division and cell death, macrophage differentiation and oxalate-driven ossification of renal epithelial cells | Hu Q. et al. 2023 |

The dot-plots derived in the study showed the activity of enriched signaling pathways to view gene clusters that are highly expressed in specific biochemical cascades (Figure 5). The enrichment dot-plots were developed using an R-package to compare the biological themes among gene clusters by interpreting the omics data and to highlight the key pathways of pathology expressed in patients with renal calcification to facilitate subsequent functional assays in the study.

**Figure 5:**
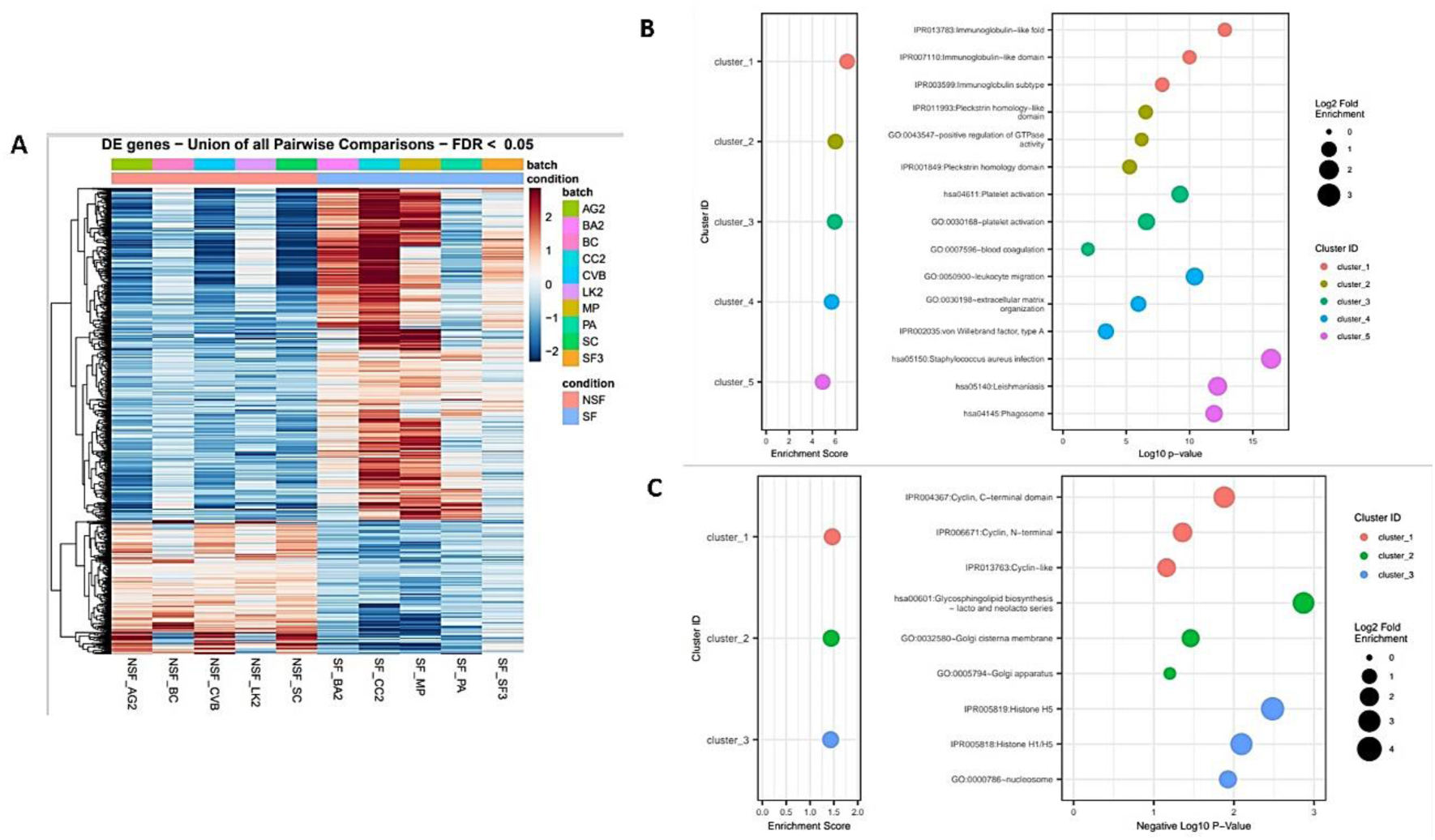
Pathway-based data integration and visualization derived from A) the heatmap generated to examine specific genes that are differentially expressed relative to inflammation, fibrosis, and calcification. B) Dot-plot enrichment assays of stone former vs. non-stone former cohorts; top-5 upregulated cluster IDs and enrichment scores, C) All upregulated clusters and their enrichment scores [Jeewandara T. et al. 2023].

The combined heatmap and dot-plot data revealed the upregulated expression of genes that are listed on table 4. For instance, the phagosome-related genes include <u>FMNL1</u>, <u>NOD2</u>, <u>RAB31</u>, <u>SLAMF1</u>and phagocytosis-promoting receptors. In the dot-plot enrichment assay, the upregulated immunoglobin-like fold correspond to genes associated with cytokine-mediated signaling such as <u>CCL19</u>, <u>CCL21</u>, <u>CCR7</u>, <u>CCR1</u>, and <u>CCR4</u>. The immunoglobin isotypes related genes mediating cell cytotoxicity and complement pathway activation include <u>FCGR2b</u>, <u>CD37</u>, <u>NIPAL3</u>, and <u>SAMHD1</u> genes. The platelet-activation gene cohort include <u>EMILIN2</u>, <u>ENTPD1</u>, <u>FERMT3</u>, <u>ITGB3</u>, and <u>HTRA1</u>. Genes related to leukocyte migration were also upregulated in the renal stone forming patient cohorts, including those associated with cellular extravasation and chemokine ligands associated with an inflammatory response, including <u>AIF1</u>, <u>CMKLR1</u>, <u>TLR8</u>, <u>NOD2</u>, and <u>SPHK1</u>. The extracellular matrix organization path upregulated in the dot-plot enrichment assay correspond to a cohort of genes, including <u>ADAMTS7</u>, <u>CCDC80</u>, <u>COL1A1</u>, <u>MMP16</u>, and <u>TGFB1</u> related to cell-matrix and collagen deposition. Zinc finger protein related genes were not observed in the heatmap of significantly differentially expressed genes, although these genes were present in the bulk-RNA sequencing data – to include <u>ZNF219</u>, <u>ZFP36</u>, <u>ZFYVE19</u>, and <u>ZFP14</u>.

**Table 4:** The combined heatmap and enrichment dot-plot derived comparisons of biological themes among gene clusters obtained via gene ontology databases, to interpret the differentially expressed genes. The analysis included all patients in the study cohort [Jeewandara T. et al. 2023].

| Enrichment pathway | List of genes | Function |
| --- | --- | --- |
| Immunoglobulin-like fold | CCL19 | Genes associated with cytokine-mediated signaling |
|  | CCL21 |  |
|  | CCR7 |  |
|  | CCR1 |  |
|  | CCR4 |  |
| Immunoglobulin-isotypes | FCGR2b | Triggering the FcγR-expressing cells for phagocytosis, or antibody-mediated cytotoxicity in cells to activate complement pathways and deliver an immune response. |
|  | CD37 |  |
|  | NIPAL3 |  |
|  | SAMHD1 |  |
| Phagosomes | FMNL1 | Phagosome related genes and phagocytosis-promoting receptors |
|  | NOD2 |  |
|  | RAB31 |  |
|  | SLAMF1 |  |
| Platelet activation | EMILIN2 | Central to homeostasis and thrombosis, also associated with renal injury and epithelial endothelial dysfunction |
|  | ENTPD1 |  |
|  | FERMT3 |  |
|  | HTR2A |  |
|  | ITGB3 |  |
| Inflammatory response | AIF1 | Genes related to leukocyte migration, including those involved in cellular extravasation and chemokine ligands associated with an inflammatory response |
|  | CMKLR1 |  |
|  | TLR8 |  |
|  | NOD2 |  |
|  | SPHK1 |  |
| Extracellular Matrix Organization | ADAMTS7 | Related to cell-matrix and collagen deposition during renal fibrosis |
|  | CCDLN80 |  |
|  | COL1A1 |  |
|  | MMP16 |  |
|  | TGFB1 |  |

### • The fluorescence in-situ hybridization (FISH) outcomes

We quantified the fluorescence in-situ hybridization outcomes in the proximal papillae and distal papillary tip regions of both stone formers vs. non-stone former patients. The outcomes showed the differential expression of transcripts between the proximal papillae and the tip region. Specific genes such as <u>GALNT3</u>, <u>PLEKHO1</u>, <u>SLCO2A1</u> and <u>VCAM1</u> were highly expressed in stone formers at the proximal papillae and not at the renal papillary tip region. The gene <u>IGF1</u> and <u>TMEM30B</u> were highly expressed at the renal papillary tip, but not at the proximal papillae. In comparison, the LOX transcript was highly expressed at the renal tip and at the proximal papillae (Figure 6).

**Figure 6:**
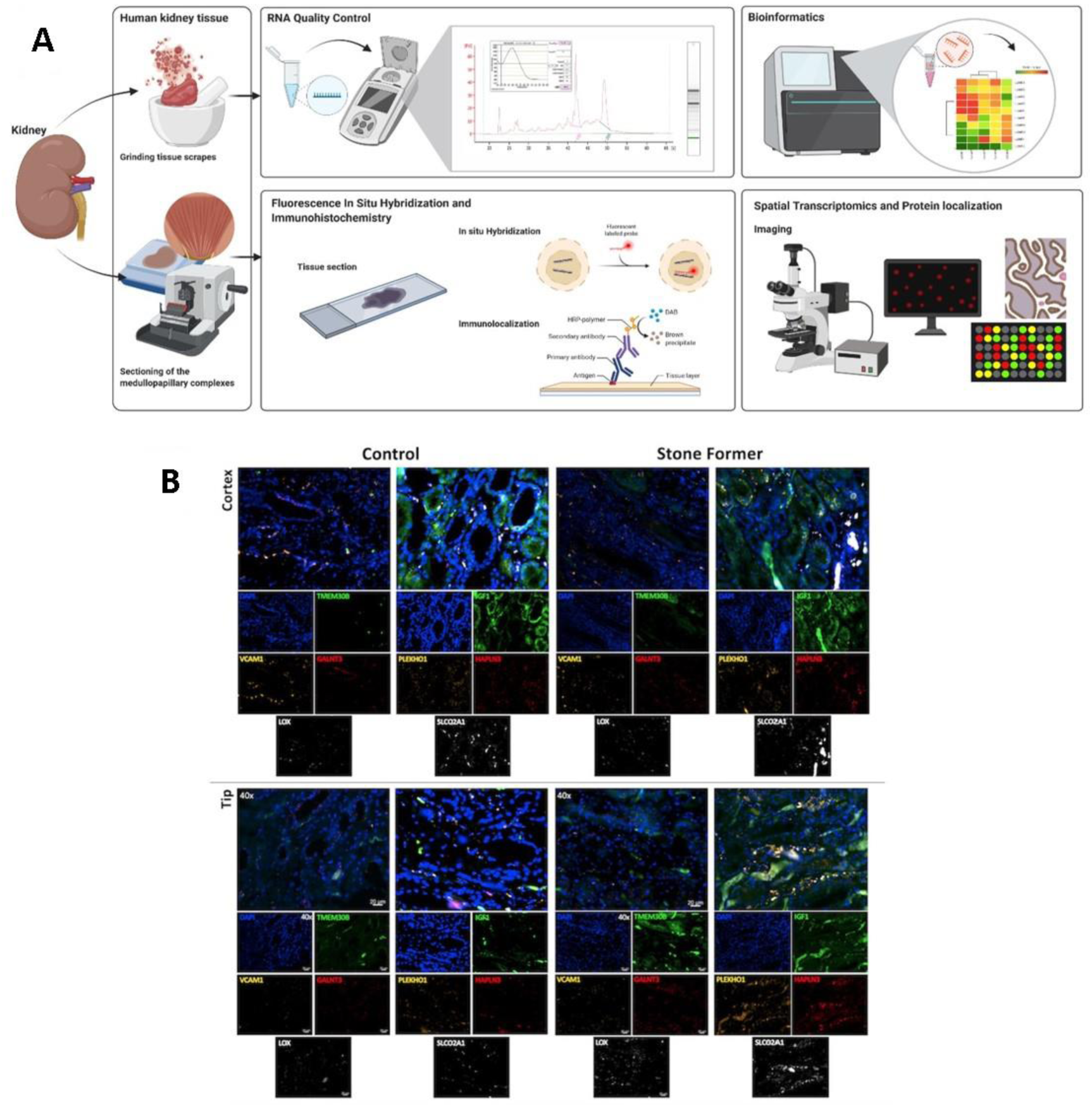
A) a schematic representation of the methods of bulk RNA sequencing and FISH conducted in-lab, B) FISH outcomes. Delineating the expression of key genes of interest (VCAM1, GALNT3, PLEKHO1, HAPLN3 and TMEM30B) in the renal papillary cortex and tip region of the control tissue specimen vs. stone formers.

### • M1 and M2 Macrophage differentiation from whole blood derived from renal stone forming patients

By day 3, the M1 and M2 macrophages derived from PBMCs had proliferated to reach confluence (Video 1). We used the macrophage detachment solution to detach the cells from the tissue culture plastic and subjected them to quality control via fluorescence-activated cell sorting (FACS). The M1 macrophages stained positive to the CD86 antibody and the M2 macrophages stained positive to the CD163 antibody to individually confirm the identification and the presence of the differentiated M1 and M2 macrophages in the cell cultures, respectively (Figure 7). Upon confirming the identity of the cells, we continued to sub-culture the cells in the static 35 mm cell culture plates and continued first-in-study macrophage cell cultures under flow conditions on the kidney-on-a-chip instrument. These cell types were continuously cultured for subsequent functional assays *in vitro*.

**Figure 7:**
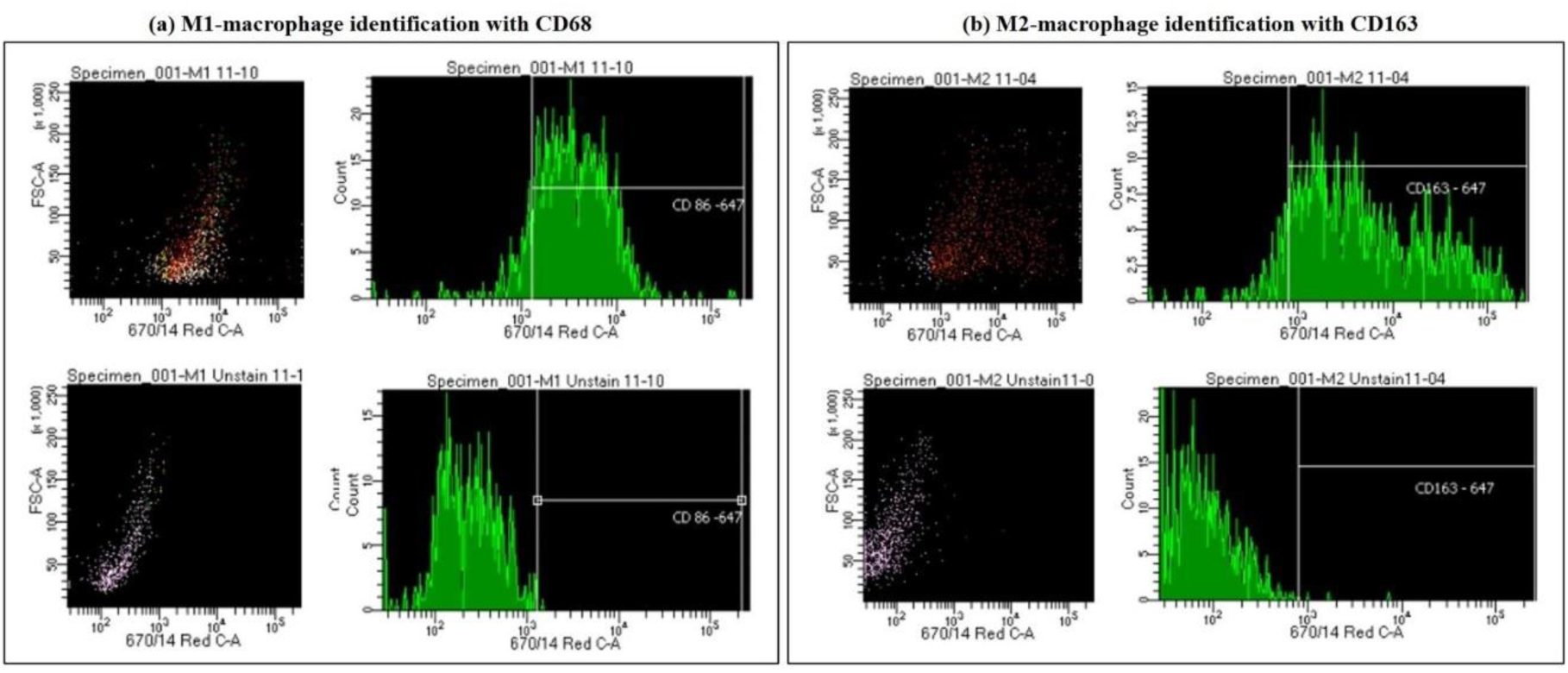
Using fluorescence-activated cell sorting (FACS) to a) distinguish M1 polarized macrophages stained with CD86 vs. the unstained M1 polarized macrophages, and b) distinguishing M2 polarized macrophages stained with CD163 vs. the unstained M2 polarized macrophages [Jeewandara T. et al. 2023].

**Video 1:** https://www.dropbox.com/scl/fi/aof4hkh56ehkw939msls5/Chip_Plate_Macrophage-Growth-Dynamics.mp4?rlkey=tidk2goyexgtuzc9ssbob97v5&st=w2d3yuwm&dl=0 a) Macrophages on a chip/plate (bright field imaging): the M1 differentiated cluster-forming macrophages have a miniature morphology and are actively motile, as they aggregate in clusters on a 35 mm plate (20 x magnification). b) Macrophages on a chip/plate (bright field imaging): the M1 differentiated cluster forming macrophages on a kidney-chip are comparatively proliferating at a higher concentration yet are well-adjusted to the material surface and grow on the organ-chip instrument at a much better rate than on a plate, showing their movement and affinity for growth at a higher density, without forming clumps or clusters (20 x magnification, highest magnification available to retain image clarity) [Jeewandara T. et al. 2023].

### • Repurposing a proximal-tubule-on-a-chip to study renal biomineralization

The Chip-S1 (Emulate Inc) was connected to a pod and to the Zoë culture hub instrument to facilitate the recreation of a versatile macrophage microenvironment with cell-cell interfaces, fluid flow and mechanical forces. The typical renal microenvironment can be created within each chip-S1 by including epithelial cells in the top channel and endothelial cells in the bottom channel, separated by a porous membrane to allow cell-cell interactions as those seen *in vivo* to emulate a proximal-tubule-on-a-chip. The two channels are fluidically independent. We repurposed the Chip-S1 to mimic the growth of renal patient-derived macrophage cell types in a microengineered environment to recreate the natural physiology and mechanical forces predominantly experienced in the medullo-papillary tip region of the kidney. We cultured the top channel with M2 macrophages and the bottom channel with M1 macrophages. The pod provided the different media to the organ-chip instrument and acted as an interface between the organ-chip and the Zoë culture module to facilitate the growth of motile, cluster-forming macrophages (Figure 8).

**Figure 8:**
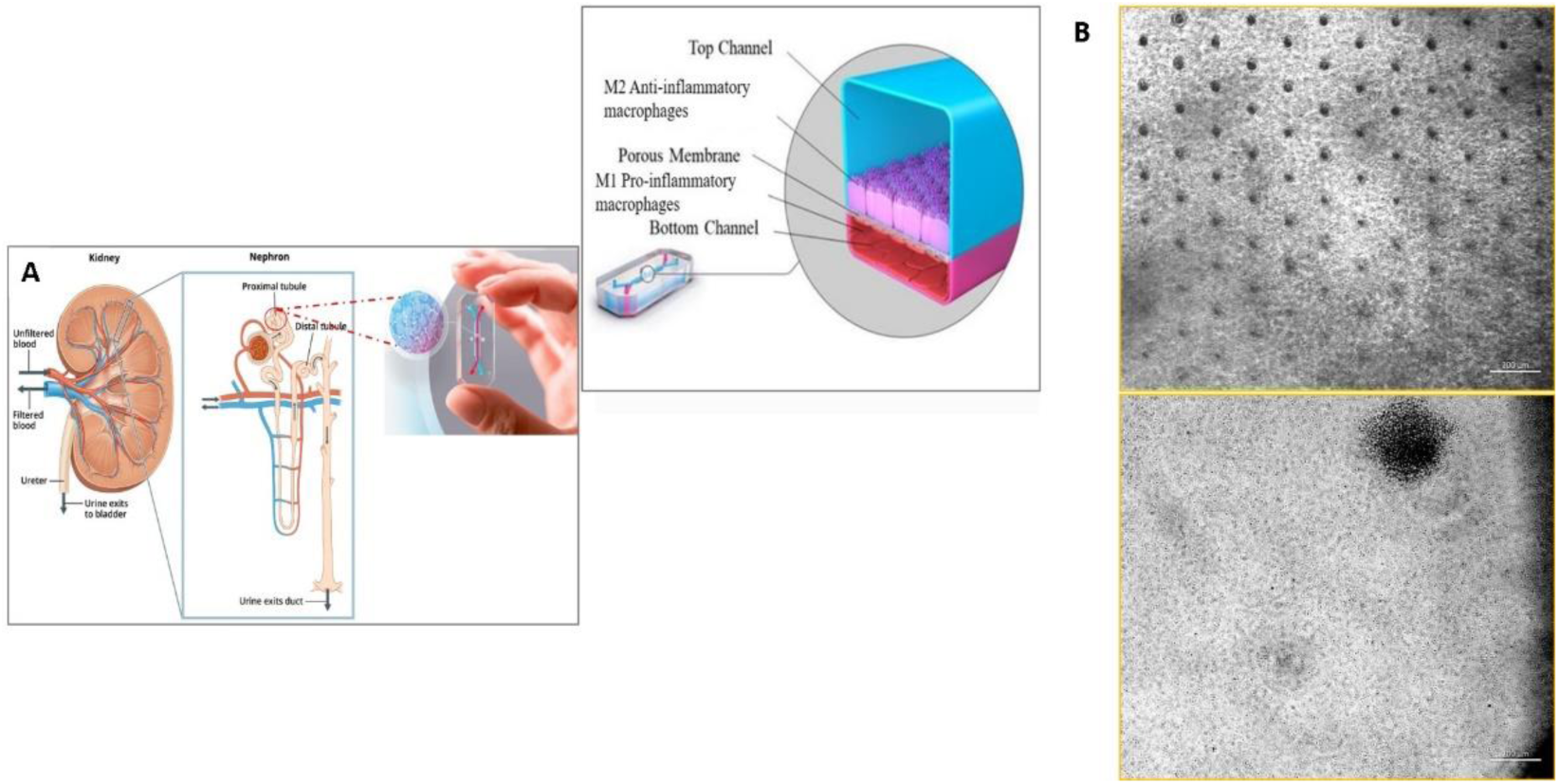
Repurposing a human proximal tubule kidney chip to A) facilitate the growth of M1 pro- and M2 anti-inflammatory macrophages. B) The bright field images are representative of cell proliferation on the kidney chip vs. a 35 mm tissue culture plate [Emulate. Inc, Jeewandara T. et al. 2023].

### • Molecular biomarkers of oxidative-stress initiated apoptosis

We subjected the macrophage cell types cultured on a kidney-chip and on a plate to apoptosis assays in the presence of sodium dithionite in oxygen glucose deprived (OGD) physiological buffer to induce oxidative stress via chemical ischemia *in vitro*. Prior to inducing apoptosis, the cell types were incubated with the CellRox redox dye (green). When we exposed cluster-forming macrophages to apoptosis assays, the cells immediately underwent aggregation, clumping, and gradual cell death. When the cell cultures were qualitatively observed under fluorescence (AG520), the cell clumps appeared to glow in green, to confirm the process of redox/oxidation *in vitro* (Figure 9) (Video 2) [<u>Jeewandara T. 2024</u>].

**Figure 9:**
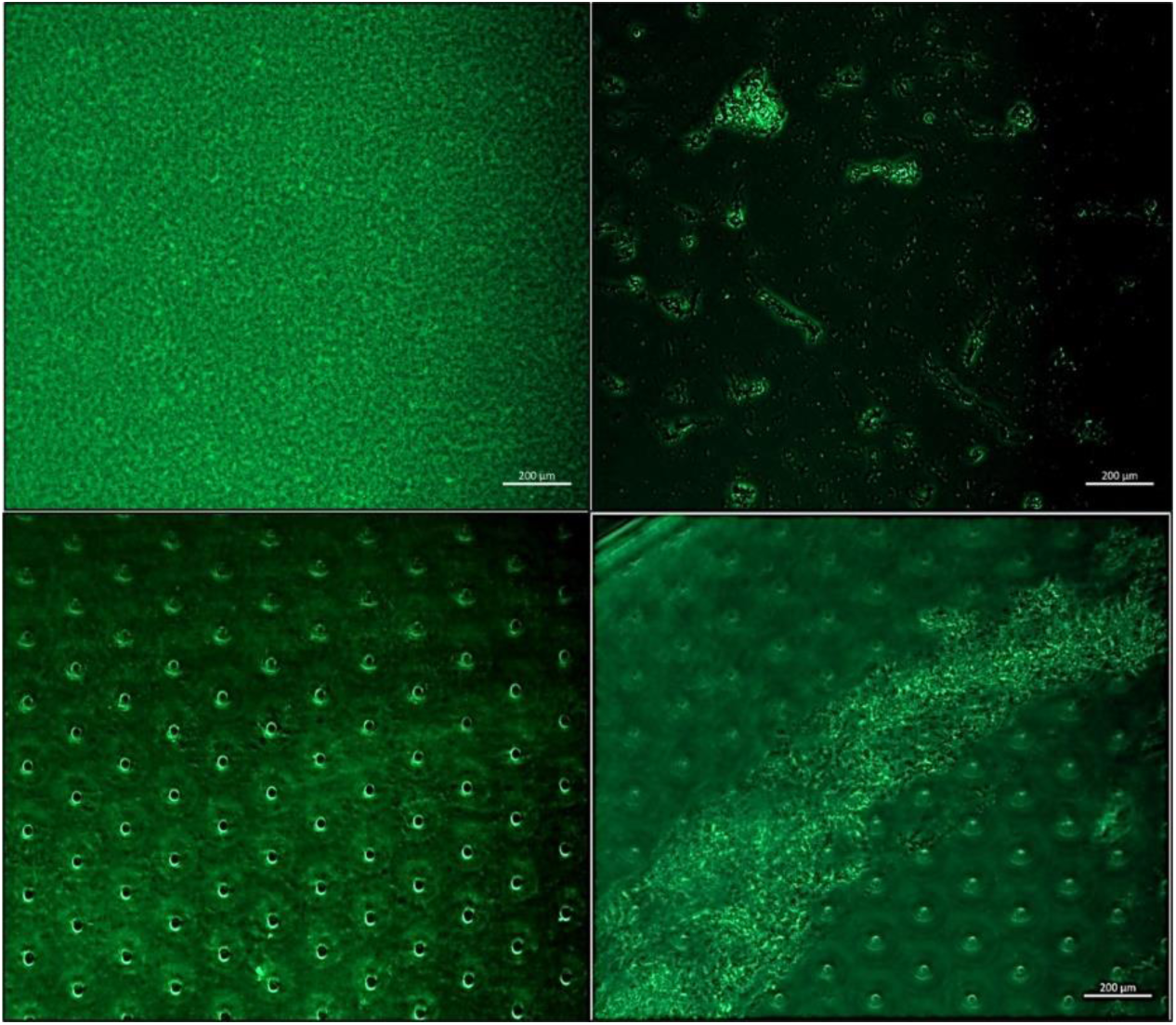
The apoptosis assays conducted with stone former M1 and M2 macrophages cultured on a plate (top) and on an organ-on-a-chip/kidney-chip instrument (bottom) incubated with CellRox (Redox) dye (green) prior to (left) - and after (right) chemically induced cell apoptosis [<u>Jeewandara T. 2024</u>].

**Video 2: Available via:** https://www.dropbox.com/scl/fi/84zlv3kiyxlezxrs9sc6f/CellRox-dye_Redox-Functional-Assay.mp4?rlkey=axp8gpe22a0hx930sch0urefs&st=einbmqlf&dl=0

Apoptosis assays on a chip/plate (fluorescence imaging): the M1 and M2 macrophage cell types were incubated with CellRox redox dye and subjected to chemical ischemia-induced redox, to observe the transition from healthy motile cells (left) to fluorescent cell clumps or aggregates (right), to indicate apoptosis or cell death under ischemia, representative culture on a 35 mm tissue culture plate. The fluorescent redox dye emission (AG520) uptake indicated oxidative stress driven cell apoptosis (20 x magnification) [<u>Jeewandara T. 2024</u>].

### • Identifying a biological switch at the onset of hypoxia-induced apoptosis with hypoxia-inducible factor 1α (HIF1α) and prolyl hydroxylase domain 2 (PHD2) interactions

Next, we subjected a fresh batch of macrophage cell cultures derived from renal patient cohorts (n=10) on a kidney-chip, and on a plate to chemically induce hypoxia, to bottom-up engineer a cell signaling pathway of hypoxia-inducible factor 1α driven oxidative stress. The omics-data revealed the HIF1α-driven hypoxia response pathway, which is primarily regulated via the key enzyme of prolyl hydroxylase domain 2 (PHD2) as a pathological switch, to mediate the hypoxia response at the early stages of renal stone formation. Prior to inducing apoptosis on the cell culture, the cells were incubated with the PHD2 antibody (orange). When the cell cultures were qualitatively observed under fluorescence (AF555), they appeared to glow in orange and emit a switch-like response as the cells underwent apoptosis under live-cell imaging (Figure 10) (Video 3).

**Figure 10:**
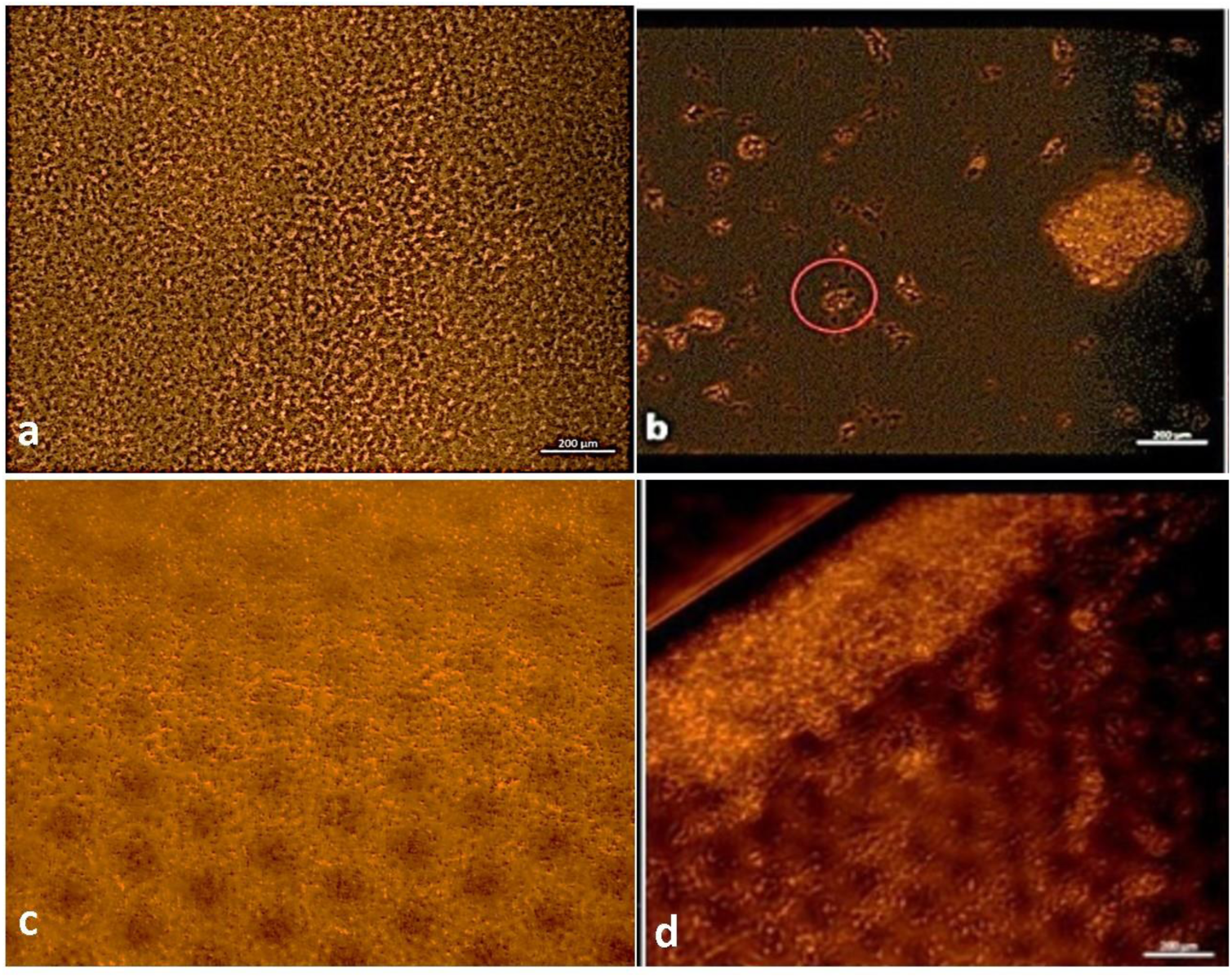
Apoptosis assays on a chip/plate (fluorescence imaging) – the M1 and M2 macrophage cell types were incubated with the AlexaFluor555 EGLN1/anti-PHD2/prolyl hydroxylase domain 2 antibody and subjected to chemical ischemia-induced redox (a-d) to observe the transition from healthy motile cells (left) to fluorescent clumps or aggregates (right) to indicate apoptosis or cell death under ischemia, represented on a 35 mm tissue culture plate and on a kidney-chip instrument. The fluorescence dye emission of AF555 represented a switch-like upregulation of PHD2 (circled in b), to glow in orange and emit a switch-like response as the cells underwent apoptosis under live-cell imaging (b,d).

**Video 3:** https://www.dropbox.com/scl/fi/stamqi8hfu39396r9t26u/Video-3_PHD2_Hypoxia-Functional-Assay_Chip-and-Plate_Annotated.mp4?rlkey=u7wvqvlan10yfbtp4kdjuftuh&st=mxyo1c8t&dl=0

Hypoxia-on-a-chip/plate (fluorescence imaging): the cell types were incubated with PHD2 (Alexa Fluor 555) antibody. Upon subjecting the M2 macrophage cells to chemically induced ischemia, they underwent oxidative stress and aggregated in immunofluorescent PHD2 expressing clumps, the activated PHD2 oxygen sensor enzyme within cells is noted as an oxidative stress signal or switch that flicker on/off, as indicated with arrows (zoom in, magnification 20 x) [Jeewandara T. et al. 2023].

## Discussion

The salient findings of this study are as follows –

1. During our first-in-study outcomes, we established a macrophage M1 and M2 cell line derived from human peripheral blood mononuclear cells (PBMCs) obtained from patients’ whole blood, and compared the expression of biomarkers of stone-former vs. non-stone former patient cohorts via appropriate functional assays.
2. We established an artificial neural network (ANN) for the quantitative pathology of biomarkers of interest in renal patient-derived papillary tissue cross-sections of histology.
3. We bottom-up engineered a pathological pathway of oxidative stress on a repurposed proximal-kidney-on-a-chip device, based on our initial omics and histopathology dataset, to recreate a pathological cell signaling pathway *in vitro*.
4. We identified the presence of a biological switch –prolyl hydroxylase domain 2 (PHD2) – a zinc finger protein enzyme that mediates the onset of hypoxia-induced factor 1α (HIF1α)-driven oxidative stress in macrophage cell samples obtained from patients, for the first time.

Our hypothesis of the presence of a biological switch that mediates oxidative stress-driven renal fibrosis and calcification at the renal papillary tip, originated from our early data of patient-derived, evidence-based, transcriptomics and histology studies conducted with clinical samples [Jeewandara T. et al. 2023]. The subsequent outcomes of our biomolecular functional assays validated our proposed hypothesis of an activated oxygen sensor enzyme, known as prolyl hydroxylase domain 2 (PHD2) that appears to function as a biological switch at the renal papillary tip in response to hypoxia. The pathological cascade originates when the hypoxia-inducible factor 1α protein is upregulated as a master transcriptional regulator during the cell response to hypoxia or oxidative stress [<u>Arsenault P. et al. 2016</u>]. We thereby established and verified a preliminary platform of oxidative stress by investigating patient-derived macrophages cultured on a repurposed proximal tubule on a chip – referred to as a kidney-chip, within the context of this paper.

As an initial study outcome, we quantified vital histopathology biomarkers that occur at the renal papillary tip region of histology tissues derived from renal stone-forming vs. non-stone forming patients. We accomplished this by programming an artificial neural network (quPath, GitHub) to measure the key biomarkers of renal calcification. Subsequently, the outcomes guided the development of our kidney-chip functional assays. For instance, the epithelial-to-mesenchymal transitions visualized in our study with vimentin-DAB are precursors of fibrosis that influence calcification, alongside macrophage-to-mesenchymal transitions [<u>Tang 2019</u>]. The biomarkers were significantly expressed in renal stone forming patient samples, when compared to non-stone formers. Collagen is another biomarker of renal injury and fibrosis visualized with Masson’s Trichrome for quantification that is typically upregulated in the renal papillary region due to shear stress [<u>Khan S. et al. 2013</u>]. Stone forming patients showed significantly higher levels of collagen deposition vs. non-stone formers. Using the Alizarin red dye, we visualized and identified renal calcification, where calcium deposits appeared as crimson aggregates across the tissue. As expected, the renal stone formers showed higher levels of calcium deposits when compared to non-stone forming patients [<u>Coralli C. et al. 2022</u>]. The papillary tip region of all stone-forming patients exhibited key biomarkers of collagen deposition (Masson’s Trichrome), epithelial-mesenchymal transitions (vimentin-DAB) and calcification (Alizarin red) in greater quantity than the non-stone formers [Jeewandara T. et al. 2023]. The histopathological outcomes and our omics-related outcomes guided our functional assays on an organ-on-a-chip device.

To initiate bottom-up bioengineering a pathological cell signaling pathway *in vitro*, we isolated peripheral blood mononuclear cells from whole blood cells obtained from stone-forming and non-stone-forming patients, as progenitor or effector cells, to give rise to the M1 pro-inflammatory and M2 anti-inflammatory macrophage cell lines. These PBMC cell types were then differentiated to form the M1 and M2 macrophage phenotypes through a proprietary protocol established in-lab [Jeewandara T. et al. 2023]. Technically, the lithogenic environments of calcification can trigger inflammation and upregulate reactive oxygen species to impair renal cells and cause M1 macrophage-mediated calcium oxalate stone deposition *in vivo* [<u>Khan S. et al. 2016</u>, <u>Khan S. et al. 2021</u>]. Typically, the M2 anti-inflammatory phenotype is associated with providing a primary defense mechanism against cellular impairments via endocytosis during early stages of renal calcification [<u>Taguchi K. et al. 2021</u>]. The M2 macrophages can clear crystal deposits from renal tissues to form an innate defense mechanism against nephrolithiasis *in vivo* (Figure 11) [<u>Khan S. et al. 2021</u>].

**Figure 11:**
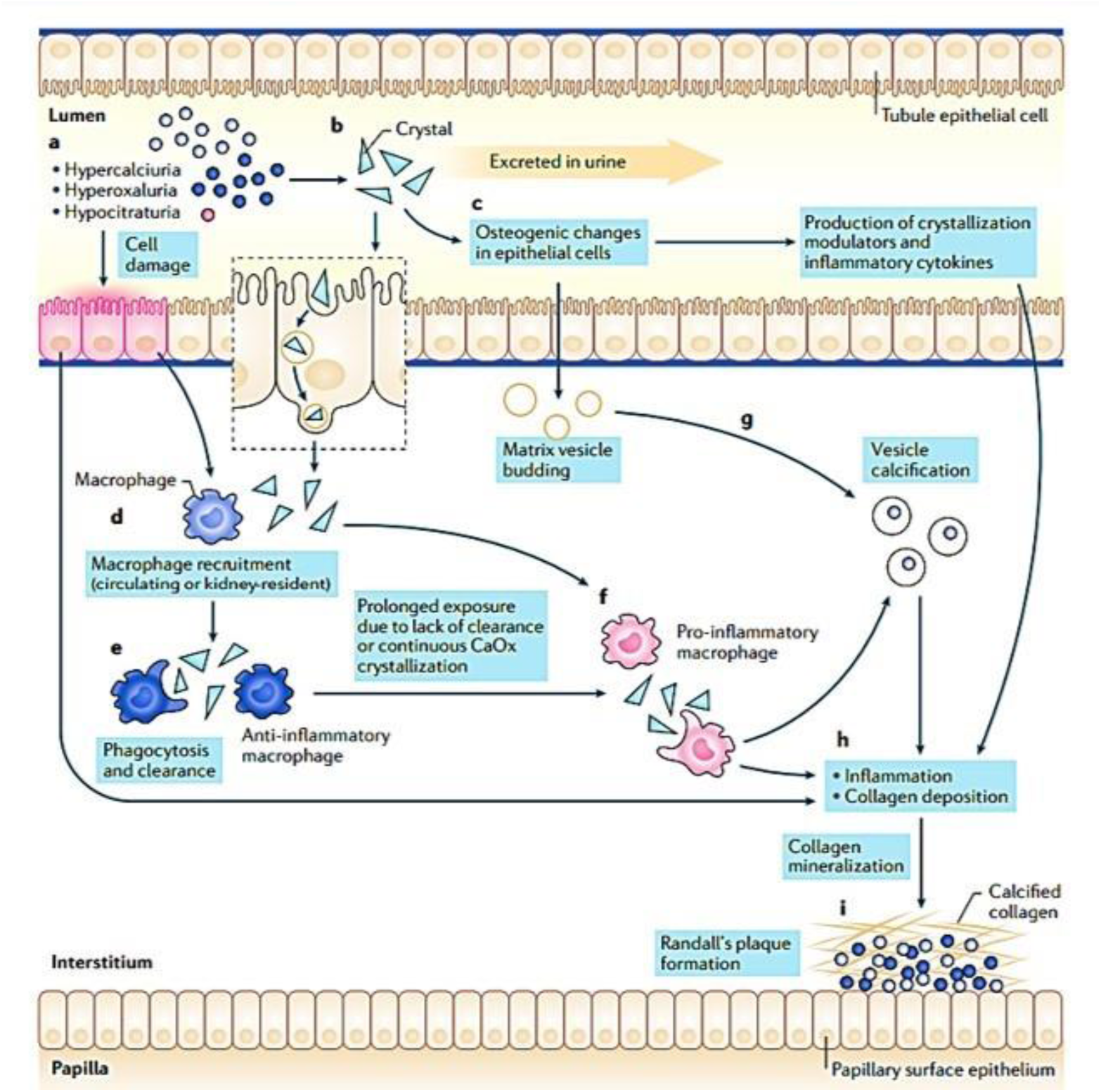
A proposed model for the formation of Randall’s plaque in the presence of M1 pro-inflammatory and M2 anti-inflammatory phenotypes. Steps a-h recap the risk factors of hyperoxaluria, hypercalciuria, and hypocitraturia and other factors involved in the pathogenesis of nephrolithiasis to cause epithelial damage, the production of calcifying vesicles, increased pro-inflammatory macrophages vs. anti-inflammatory macrophages, collagen deposition, mineralization, and Randall’s plaque formation [<u>Khan S. et al. 2021</u>].

Macrophages play a significant role during the early stages of hyperoxaluria-directed renal calcification known as Randall’s plaque formation [<u>Tang 2019</u>]. During its proposed mechanism-of-action at nephrolithiasis, M2 anti-inflammatory phenotypes ingest particulate materials to clear circulating calcium oxalate crystal deposits from renal papillary tip regions. Comparatively, the M1 macrophages initiate collagen deposition, kidney-injury mediated epithelial damage and endothelial dysfunction for pathological biomineralization, including Randall’s plaque formation [<u>Khan S. et al. 2021</u>]. First reported in 1999, this avenue can be clinically explored to offer a solution by developing treatments for kidney stones [<u>de Water 1999</u>]. To accomplish this, it is necessary to understand the nature of pathological cascades that trigger the diverse roles of macrophages during renal crystal origin, prompting the cells to migrate to calculi deposition sites and to engulf the crystals [<u>Murray P. et al. 2014</u>]. These observations prompted our investigations on the M1 and M2 macrophage cell lines in our study.

Our study outcomes further demonstrated the versatility of the kidney-on-a-chip instruments. We observed better support and proliferation of the miniature, cluster-forming motile M1 and M2 macrophages on the kidney-chip instruments, when compared to similar macrophage aggregates that differentiated into clusters on the 35 mm tissue culture plate surfaces. The extracellular matrix of the organ-chip instrument is designed on the first-principles of biology to support cellular ‘tensional integrity,’ by mimicking cell and tissue adhesion *in vivo* [<u>Ingber D.E. 1993</u>]. The relatively simple theory, underlies much of the complexity of pattern and structure within the cytoskeleton of living cells to facilitate cell proliferation and cellular organization that contributes to information processing, mechanochemical transduction and morphogenetic regulation [<u>Ingber D. E. et al. 2015</u>], which we emulated on the instrument *in vitro*. In our study, we further highlighted the possibility of repurposing kidney-on-a-chip instruments without using an artificial membrane to simulate the filtration membrane of the filtering kidney, by recreating parameters of pathological shear-stress *in vitro*. We developed an extracellular matrix using collagen and Matrigel proteins to facilitate cell attachment and proliferation within the renal microenvironments based on first-principles of tensegrity to effectively recreate microphysiological conditions on a kidney-on-a-chip instrument, and bottom-up engineer a pathological cell signaling pathway to conduct functional assays *in vitro*. Next, we identified the macrophages differentiated in the lab via FACS to distinguish between the two types of polarizations. Biologically, the M2-like macrophages have greater capacity to phagocytose crystals (anti-inflammatory role) than M1-like macrophages (pro-inflammatory role) that are instead involved in the process of crystal deposition. The capacity to regulate M2 polarization therefore has therapeutic value [<u>Taguchi K. et al. 2021</u>].

Based on the transcriptome analysis of our study, we used the Panther database and DAVID-KEGG to detect a spectrum of 64 pathological pathways that are predominant to renal stone forming patients, of which we selectively focused on the hypoxia-inducible factor activation (HIF1α) pathway as a primary pathological cascade of interest, due to its influence on renal calcification [<u>Negri A. 2023</u>]. Of the transcriptome data, we highlighted the differentially expressed genes that were significantly upregulated in association with phagocytosis, cytokine mediated signaling, and complement pathway activation to deliver an immune response. We further observed genes associated with cell-matrix and collagen deposition during renal fibrosis that are linked to renal stone-forming patients (Table 4). The transcriptome analysis demonstrated cyclin-like, cyclin-c-terminal and cyclin-n-terminal domains that regulate the cell cycle associated with renal carcinoma of the ‘non-stone former’ patient cohort [<u>Ghafouri-Fard S. et al. 2022</u>, <u>Sukov W. et al. 2009</u>]. The outcomes also highlighted the upregulation of histone deacetylases associated with the epigenetics machinery underlying acute kidney injury, with scope to develop histone deacetylase inhibitors as small-molecule drugs to treat kidney injury associated with renal calcification [<u>Hyndman 2020</u>]. Our observation of the upregulation of a zinc finger protein enzyme PHD2 as an oxygen sensor at the onset of chronic hypoxia response-related calcification, coincides with our transcriptome analysis data that also showed the abundant expression of Zinc finger proteins in tissue samples of patients. Future technical investigations of the elemental composition of the renal papillary tip region of clinical stone-formers via methods in materials science such as energy dispersive x-ray spectroscopy (EDX) can delineate the expression level of the zinc signal at the onset of hypoxia, and determine if the signal progressively decreases with time during progressive calcification of nephrolithiasis.

The ‘hypoxia response via HIF activation’ pathway is often upregulated in the transcriptome of stone forming patients and is a key mechanism of HIF1α stabilization [<u>Han W. et al. 2013</u>]. In its mechanism-of-action, prolyl hydroxylase domain 2 (PHD2) protein is a primary oxygen sensor/regulator and a site-specific zinc finger enzyme that can hydroxylate HIF1α and degrade it under normoxia by forming a complex with the von-Hippel-Lindau factor (pVHL) [<u>Arsenault P. et al. 2016</u>]. Under hypoxia, this process of degradation does not occur, leading to the stabilization of HIF1α instead, facilitating its subsequent activation as a pathological hallmark of kidney injury and stone formation (Figure 12) [<u>Meneses 2016, Arsenault 2016</u>]. During the experiments, we induced chemical-ischemia mediated cell death via apoptotic assays in the presence of renal patient-derived M1 pro and M2 anti-inflammatory macrophage cell lines cultured on a microfluidic kidney-on-a-chip instrument and on a culture plate *in vitro*. Our first-in-study outcomes emulated pathological conditions on a kidney-chip and a plate to verify the biological switch-like signaling and activation of PHD2 during oxidative stress initiated at the onset of ‘hypoxia response via HIF activation.’

**Figure 12:**
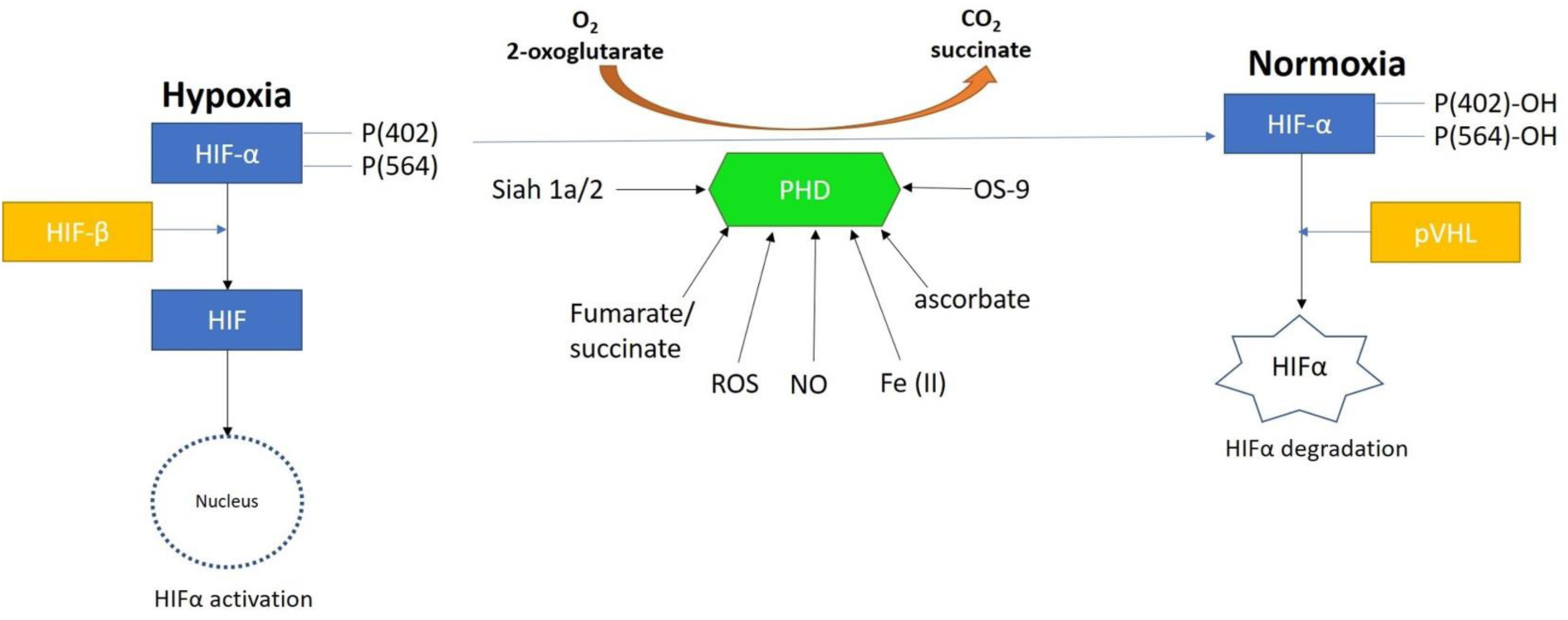
Prolyl hydroxylase domain 2 (PHD2) is a key oxygen sensor in mammals that post-translationally modifies hypoxia-inducible factor α (HIF1α) to target it for degradation by forming a complex with the Von Hippel-Lindau factor (pVHL). Reactive oxygen species, nitrous oxide, osteosarcoma amplified 9, ascorbate, succinate, iron, and protein ligase components can regulate the function of PHDs [recreated via <u>Arsenault P. et al. 2016</u>].

We studied the hypoxia-induced oxidative stress pathway, since the hypoxia inducible factor 1α (HIF1α) protein is associated with progressive oxidative stress-related chronic renal injuries, tissue fibrosis, and the genetic expression of vimentin – a biomarker of epithelial-to-mesenchymal transitions and cell death [<u>Higgins D. et al. 2007</u>]. These biomarkers were noted at the onset of our histology experiments too – where the expression of vimentin, collagen and calcium deposits were significant at the renal papillary tip of clinical stone-former patient samples. These pathological biomarkers subsequently guided our functional assays developed on a chip and on a tissue culture plate – to bottom-up engineer the pathological cell signaling cascade of interest, *in vitro* [Jeewandara T. et al. 2023].

Recreating or emulating oxidative stress on an organ-chip instrument deals with the premise that reactive oxygen species are key regulators of disease and unite many mechanisms for calcium oxalate stone formation *in vivo* [<u>Khan S. et al. 2014</u>]. Tissue culture studies that were previously conducted with renal epithelial cells exposed to high concentrations of oxalate, calcium oxalate or calcium phosphate crystals have caused injury to cell types [<u>Khan S. et al. 2014</u>, <u>Koul H. et al. 1994</u>]. Such cell exposure can cause the secretion of superoxide in an extracellular oxidative burst. The resulting cellular injury can be ameliorated in-lab with antioxidants and free radical scavengers such as catalase, and superoxide dismutase to protect cells from oxalate-induced injury [<u>Gaspar S. et al. 2010</u>, <u>Thamilselvan S. et al. 2000</u>]. Additionally, the interaction of calcium oxalate monohydrate crystals with renal cells can cause altered gene expression, and initiate DNA synthesis as well as cell death, factors that are mediated by the p38 MAPK pathway to activate macrophage-facilitated inflammation (Figure 13) [<u>Koul H. et al. 2002</u>, <u>Hammes M. et al. 1995</u>].

**Figure 13:**
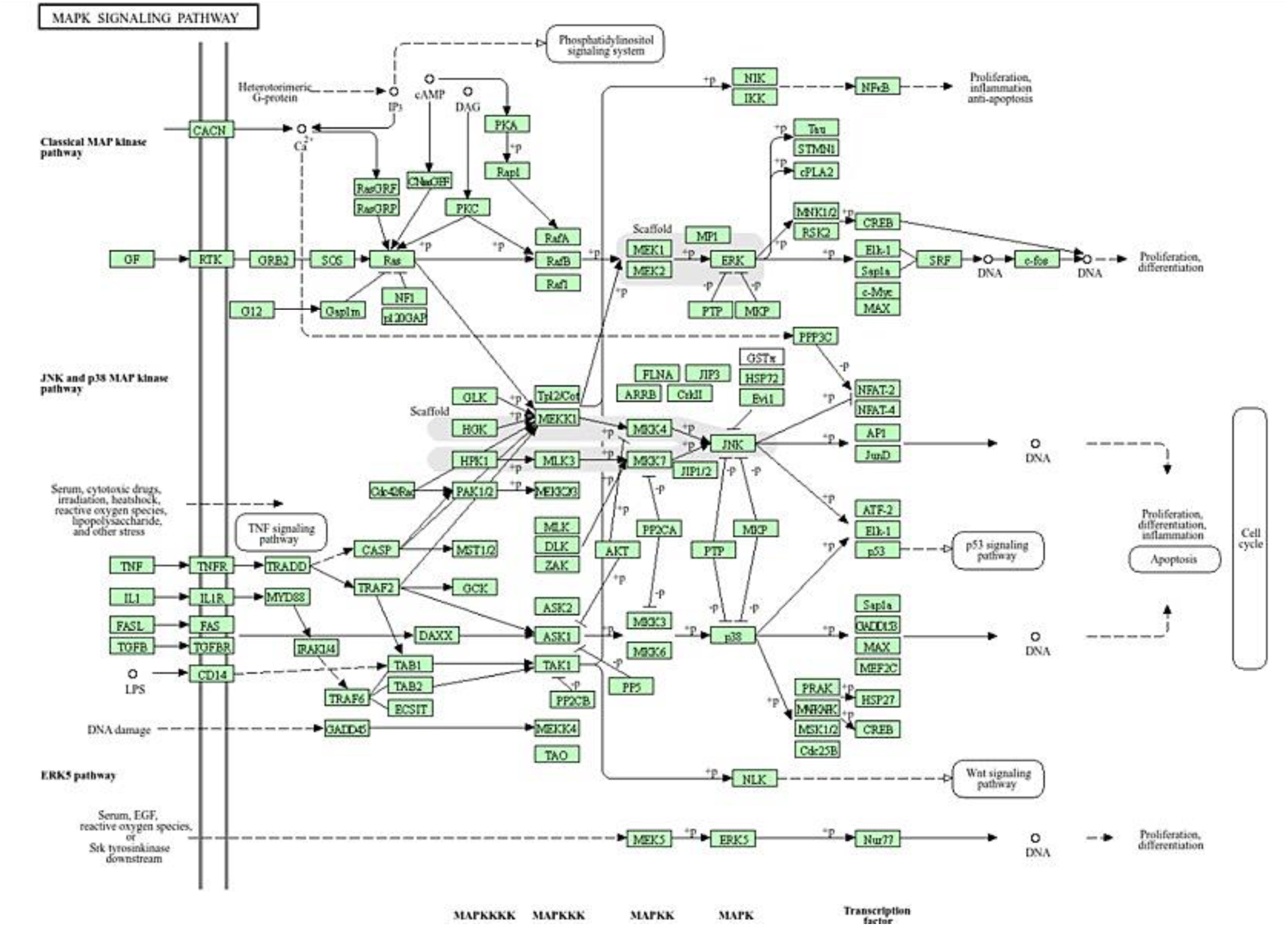
The KEGG metadata pathway generated of the patient-derived transcriptome-driven bioinformatics data to elucidate key cascades of renal calcification – depicting the p38 mitogen activated protein kinase (p38 MAPK) pathway associated with cell stress and macrophage activation to mediate an inflammatory response [<u>KEGG</u> <u>database</u>].

Preclinical research in animal models of calcium oxalate nephrolithiasis and hyperoxaluria have thus far shown that the treatment with antioxidants such as vitamin E improved the tissue levels of antioxidant enzymes, to reduce injury and eliminate calcium oxalate crystal deposition in the kidneys [<u>Thamilselvan S. et al. 2005</u>]. For example, the administration of an antioxidant apocynin can nearly completely reverse the effects of hyperoxaluria in animal models, and reduce the deposition of calcium oxalate crystals in kidneys as a proof-of-concept [<u>Khan S. et al. 2014</u>]. Such functional assays can be replicated with clinical samples of hyperoxaluric epithelial and endothelial cell types derived from renal stone former biopsies to ultimately observe the therapeutic effects of administering small molecule antioxidant drugs on a chip *in vitro*.

Since the discovery of oxygen sensors in 1996 revealed the mechanistic basis for the cellular response to hypoxia, and paved the way to therapeutically target the response to treat several diseases including cancer and anemia [<u>Forsythe J. et al. 1996</u>]. Our preliminary investigations of hypoxia-induced nephrolithiasis can be supported with further research works to identify its genetic basis. Starting with <u>egl-9-family hypoxia inducible factor 1</u> (EGLN1) that encodes the biological switch-like prolyl hydroxylase domain 2 (PHD2) enzyme driven response pathway that is underlying the onset of pathological biomineralization. Further investigations can also examine the trajectory influencing the mechanosensitive ion channels such as Piezo1 and the transient receptor potential vanilloid subfamily 4 (TRPV4) (<u>Coste B. et al. 2010</u>) that contribute to shear-stress-driven renal fibrosis that precedes renal calcification or kidney stone formation. During its mechanism-of-action, shear-stress related increases in mechanotransduction can convert physical forces of stress into biochemical signals to trigger tissue injury that eventually leads to progressive renal fibrosis and calcification (<u>He et al. 2022</u>).

Our pathological cascade of interest in the study, the ‘hypoxia response via HIF activation’ pathway was exclusively investigated on repurposed kidney-on-a-chip platforms with macrophage cell lines. Additional outcomes of our transcriptomics data for stone-former vs. non-stone former patient-derived RNA sequencing, revealed a variety of pathological cell-signaling pathways, such as the phagosomes pathway and the FcγR-mediated phagocytosis pathway to highlight the role of professional phagocytic macrophages during kidney stone formation (Figure 14). The p38 MAPK pathway represented yet another avenue of the macrophage-mediated inflammatory response that can be harnessed for therapeutic investigation [<u>Yang Y. et al. 2014</u>, <u>Raza A. et al. 2017</u>] to substantiate our research focus on macrophages as versatile players in renal inflammation and fibrosis [<u>Tang 2019</u>].

**Figure 14:**
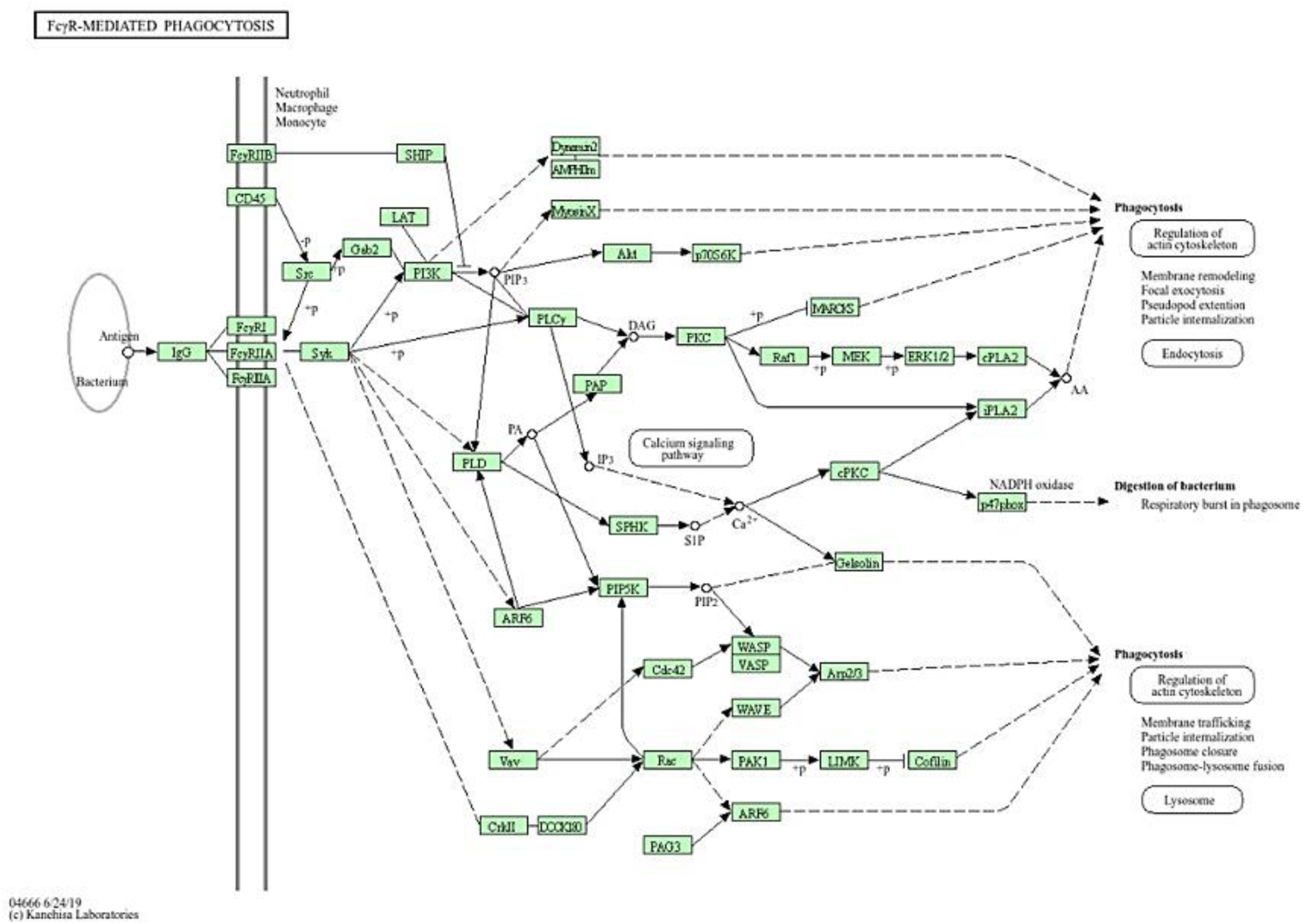
The KEGG metadata pathway generated of the patient-derived transcriptome-driven bioinformatics data to elucidate key cascades of renal calcification – depicting the FcγR-mediated phagocytosis pathway. Associated with macrophage cell surface immune receptors to clear calcium deposits during renal calcification [<u>KEGG database</u>].

Besides transcriptomics, we conducted fluorescence in situ hybridization (FISH) studies to identify a select set of genes including biomarkers of inflammation by using patient-derived medullopapillary complexes. The study outcomes notably highlighted the heightened expression of the endothelial cell adhesion molecule – vascular adhesion molecule-1 (VCAM1) as a key mediator of endothelial dysfunction. The protein VCAM-1 can be induced via several factors that accompany inflammation *in vivo*, such as reactive oxygen species, cytokines, and turbulent stress [<u>Cook-Mills J. et al. 2011</u>]. Shear stress can regulate these biochemical signals that are involved in downstream signaling, to trigger endothelial dysfunction and fibrosis [<u>Chiu J. et al. 2003</u>]. Tissue fibrosis is a precursor of disease and an intermediate clinical feature seen at the onset of calculi formation, and urolithiasis. The capacity of shear-stress related mechanotransduction to convert physical forces into biochemical signals and trigger tissue injury and subsequent fibrosis-related renal calcification is another key pathological trajectory that can be investigated on a kidney-on-a-chip instrument.

The epigenetic cellular machinery underlying the HIF1α-driven oxidative stress pathway can be investigated for deeper therapeutic insights (Figure 15), to translate to a kidney-chip precision healthcare platform. During chronic hypoxia for instance, when oxygen sensor molecules such as PHD2 are effectively inhibited, the pathological trajectory upregulates the lysine methyl transferase enzymes G9a and G9a-like proteins (GLP), which then directly bind to the alpha subunit of HIF1α to induce methylation for continued inhibition of the hypoxia protein [<u>Chopra A et al. 2020</u>]. In its mechanism-of-action, DNA methylation can be mediated via G9a and G9a-like proteins to suppress HIF1α transcription activity and inhibit HIF1α, by reducing its transactivation domain function [<u>Shinkai Y. 2011</u>]. This is an alternate trajectory of HIF1α degradation, when compared to the PHD2-regulated pathway of von Hippel Lindau protein binding and ubiquitination of HIF1α [<u>Arsenault P. et al. 2016</u>].

**Figure 15:**
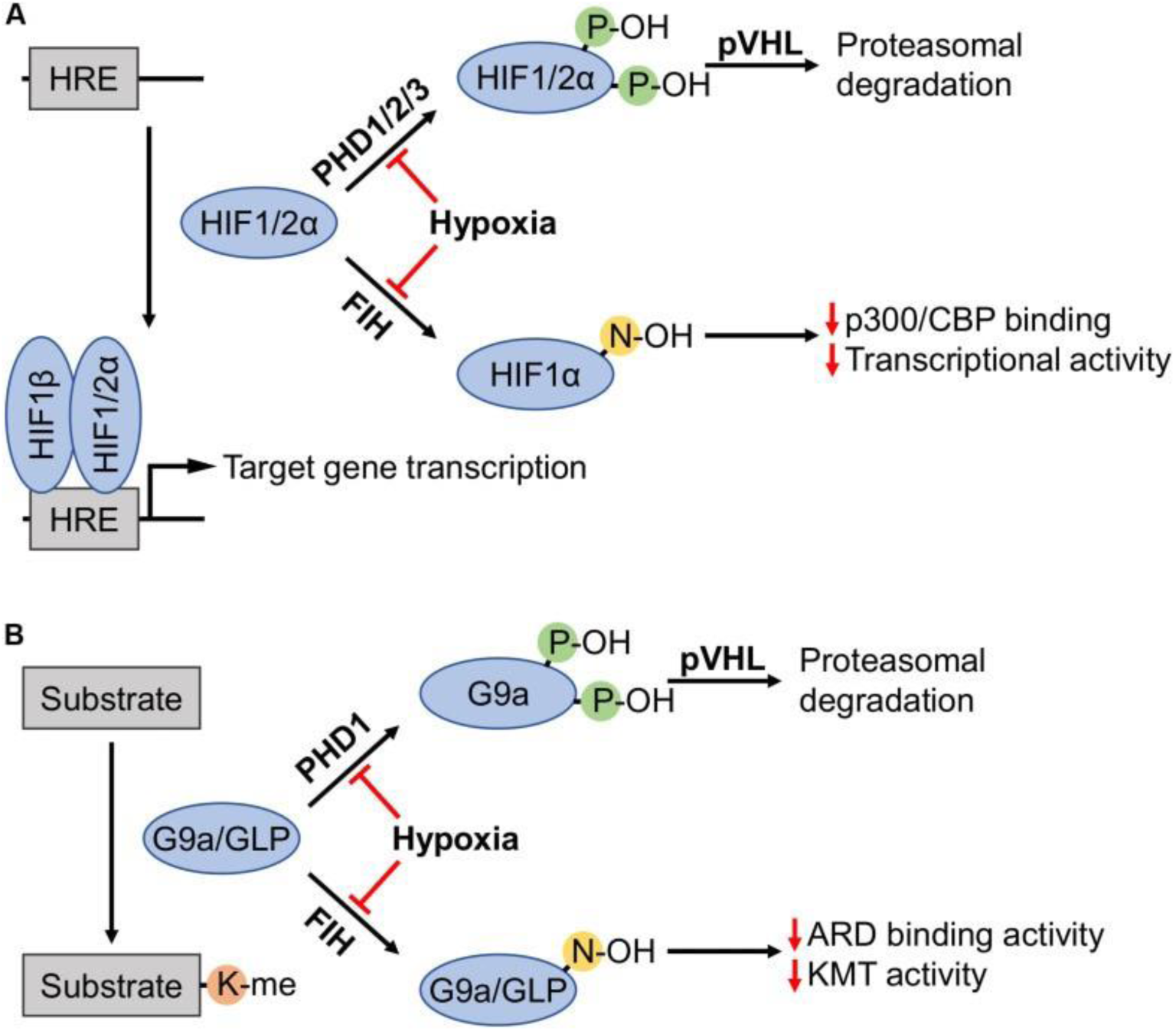
Highlighting the proteasomal degradation pathway and the regulation of HIF1α, G9a and GLP by the oxygen sensor prolyl hydroxylase domain 2 enzyme (PHD2) under normoxia. (A) Prolyl hydroxylases (PHDs) in the presence of sufficient oxygen levels (normoxia) induce the recognition and degradation of HIF1α via von Hippel Lindau tumor suppressor protein (pVHL). Factor inhibiting hypoxia has a similar role. (B) At the onset of hypoxia, HIF1α undergoes methylation induced via G9a and GLP for its suppression as an alternate pathway of HIF1α degradation [<u>Chopra 2020</u>].

This alternate path highlights the epigenetic role of G9a to be larger than what is currently known [<u>Yokoyama M. et al. 2017</u>]. This avenue can provide the G9a compound greater scope as an existing druggable target to design personalized molecular medicine, and regulate pathological hypoxia in patients pre-disposed to chronic renal calcification. At normoxia, the oxygen sensor molecule PHD2 can suppress all involved factors of the epigenetic pipeline; including HIF1α, the hypoxia-inducible G9a, and the GLP proteins, to attenuate oxidative stress-related factors and reach equilibrium. The methylation-driven epigenetic mechanism can be investigated during oxidative stress-related renal biomineralization, with cross-disciplinary impact to also regulate oncogenesis [<u>Menses A. et al. 2016</u>, <u>Yokoyama M. et al. 2017</u>].

In this way, all complex diseases result from a concerted effort of several dysfunctional pathways that come together to establish disorder – the pathology of calcification follows a similar trajectory [<u>Khan S. et al. 2021</u>]. We have already shown how the bulk RNA-sequencing data can be analyzed from renal papillary tip regions of stone forming patients to reveal the genetic upregulation of several pathological cell-signaling biomechanisms, during renal calculi formation [Jeewandara T. et al. 2023]. These gene analyses broadly include, actin cytoskeleton dysregulation, the upregulation of phagosomes, hypoxia response via HIF1α activation, apoptosis signaling pathways, p38 MAPK pathway activation and oxidative stress [<u>Koul H. et al. 2002</u>]. Each of these pathological cascades can be similarly independently investigated on a kidney-chip instrument to understand the key cell signaling pathways associated with nephrolithiasis. In agreement with our transcriptomics study outcomes, the first-principles based functional assays substantiated the presence of pathological biomarkers of oxidative stress, F-actin dysregulation, and macrophage infiltration that occur at the renal papillary tip region during calculi formation. Our preliminary and first-in-study experimental outcomes highlighted the biological switch-like activation of the prolyhydroxylase domain 2 (PHD2) enzymes to regulate the hypoxia inducible factor 1α (HIF1α) protein, and activate a route of oxidative stress in the renal papillae of stone-forming patients. Our initial experimental capacity to reveal a switch-like regulator of pathological oxidative stress is a significant observation, with scope for eventual therapeutic intervention in the clinical treatment of nephrolithiasis by forming a precision healthcare directive to treat kidney stone formers.

## Conclusion

Our first-in-study outcomes have shown the presence of prolyhydroxylase domain 2 (PHD2) as a pathological switch that can regulate the hypoxia inducible factor 1α (HIF1α) protein in the renal papillary tip region during normoxia, and at the onset of hypoxia induced on renal patient-derived tissues that were cultured on a repurposed kidney-on-a-chip instrument. We bottom-up engineered the pathological cell signaling pathway of hypoxia response via HIF activation due to its clinical relevance, substantiated with histopathology and transcriptomics data in our study. To conduct the functional assays of apoptosis induced under chemical ischemia, we established and verified healthy macrophage M1 and M2 cell lines in the lab for the first time via a proprietary protocol. Our functional assays were guided by our preceding experimental investigations on transcriptomics (bulk-RNA sequencing) and genomics (FISH) studies that highlighted the expression of cell signaling pathways including hypoxia response via HIF activation, p38 MAPK pathway, actin cytoskeleton regulation, and FcγR-mediated phagocytosis cascades.

The genomics outcomes also highlighted the pathological expression of VCAM-1 associated with mediating endothelial dysfunction *in vivo*, and the expression of biomarkers of early renal fibrosis and calcification including collagen, calcium, vimentin, as well as epithelial-mesenchymal transitions. In our work, we also developed a proprietary artificial neural network to automate the classification and quantification of histology sections derived from the renal papillary tip region of stone-forming and non-stone forming patients, to identify biomarkers of interest in tissue sections collagen, calcium, vimentin, as well as epithelial-mesenchymal transition, similar to those observed with transcriptomics analysis, to highlight the consistency of the pathological biomarkers of renal fibrosis and endothelial dysregulation that accompany nephrolithiasis. In this work, we highlighted a single pathological cascade of ‘hypoxia response via HIF activation’ that underlies renal calcification and investigated it with appropriate functional assays. The study outcomes encourage further investigations of interlinked pathological trajectories that drive renal calcification to be similarly studied on an organ-on-a-chip instrument *in vitro*. Additional pre-clinical studies can also focus on inhibiting the renal calcification trajectory by introducing small molecule therapeutic drugs-on-a-chip that are designed to combat oxidative stress and facilitate its epigenetic regulation, for translational value.

## Acknowledgements

The abstract of this manuscript was originally produced and accepted as a document submission to the scientific sessions of the World Congress of Nephrology 2023, via the University of California San Francisco (UCSF), titled “*The dynamics of pathological biomineralization in the renal papillae*.” Abstract number: WCN23-0323 in December 2022/January 2023. The initial study outcomes and future directions of the study were published on Springer Nature Research Communities, Nature Portfolio. The following publications are accessible; *“Bottom-up engineering a pathological cell signaling pathway on an organ-on-a-chip instrument”* (<u>Dec 01,</u> <u>2022</u>), *“The influence of mechano-active ion channels during kidney stone formation”* (<u>May 01,</u> <u>2024</u>), *“Experimental biology – bioinspired engineering on a kidney chip to investigate renal calcification”* (<u>July 20, 2024</u>), and *“Precision Health on a kidney-on-a-chip: developing state-of-the-art microphysiological platforms”* (<u>August 20, 2024</u>). The author acknowledges the following:

The UCSF Medical School with clinical collaborations with the Departments of Urology and the UCSF Medical Center for the clinical sample procurement and for the cell culture laboratories.

The Zuckerberg San Francisco General Hospital (SFGH) for the histopathology-immunohistochemistry collaborations.

Emulate.Inc for the organ-on-a-chip instruments, bio-kits, reagents, Matrigel/collagen compounds and for organizing the Emulate Training Agenda for the initial in-lab training sessions at UCSF.

Members of the Biological Imaging Development co-labs of UCSF for assisting the development and optimization of the artificial neural network-based quPath software.

Members of the Parnassus Flow Cytometry CoLab unit at UCSF for their assistance with quality assurance and FACS quality assessment to identify M1 and M2 macrophage phenotypes in lab.

Interdisciplinary collaborators in the Departments of Urology, Preventative and Restorative Dental Sciences, and Orthopedics in the Schools of Medicine and Dentistry at UCSF and from UC Berkeley.

This study was funded by the University of California San Francisco, California 94143.

## References

1. Levey A. and Coresh J., Chronic kidney disease, Lancet 2012

2. Wiener S. et al. Novel Insights into Renal Mineralization and Stone Formation through Advanced Imaging Modalities, Connective Tissue Research 2019

3. Ho S. et al. Architecture-Guided Fluid Flow Directs Renal Biomineralization, Scientific Reports, 2018

4. Coe F. et al. Three pathways for human kidney stone formation, Urological Research, 2010

5. Jeewandara T. et al. The dynamics of pathological biomineralization in the renal papillae, World Congress of Nephrology 2023, abstract number: WCN23:0323, 2023.

6. Rao C. et al. Effects of physical properties of nano-sized hydroxyapatite crystals on cellular toxicity in renal epithelial cells, Materials Science and Engineering: C, 2019.

7. Evan A. et al. Mechanisms of human kidney stone formation, Urolithiasis, 2014

8. Jeewandara T. et al. Experimental biology – Bioinspired engineering on a chip to investigate renal calcification, SpringerNature Research Communities, 2024

9. Saenz-Medina J. et al. Endothelial Dysfunction: An Intermediate Clinical Feature between Urolithiasis and Cardiovascular Diseases, International Journal of Molecular Sciences, 2022

10. Nature Collection: Nobel Prize in Physiology or Medicine 2019

11. Forsythe J. et al. Activation of vascular endothelial growth factor gene transcription by hypoxia-inducible factor 1, Molecular Cell Biology, 1996

12. Schofield C. and Ratcliffe P., Oxygen sensing by HIF hydroxylases, Nature Reviews Molecular Cell Biology, 2004

13. Chopra A et al. Hypoxia-Inducible Lysine Methyltransferases: G9a and GLP Hypoxic Regulation, Non-histone Substrate Modification, and Pathological Relevance, Frontiers in Genetics, 2020

14. Ingber D. E. Is it Time for Reviewer 3 to Request Human Organ Chip Experiments Instead of Animal Validation Studies? Advanced Science, 2020

15. Ingber D.E. Cellular tensegrity: defining new rules of biological design that govern the cytoskeleton, Journal of Cell Science, 1993

16. Arbra C. et al. Microdissection of Primary Renal Tissue Segments and Incorporation with Novel Scaffold-free Construct Technology, Journal of Video Experiments, 2018.

17. Bankhead P. et al. QuPath: Open-source software for digital pathology image analysis, Scientific Reports, 2017

18. Ashburner M. et al. Gene ontology: tool for the unification of biology. The Gene Ontology Consortium, Nature Genetics, 2000

19. Mi H et al. Large-scale gene function analysis with the PANTHER classification system, Nature Protocols, 2013

20. Mi H. et al. Protocol Update for large-scale genome and gene function analysis with the PANTHER classification system (v.14.0), Nature Protocols, 2019

21. Yu G. et al. clusterProfiler: an R Package for Comparing Biological Themes Among Gene Clusters, OMICS, 2012.

22. Otsu N., et al. A Threshold Selection Method from Gray-Level Histograms, IEEE Xplore 1979.

23. Nookala A. et al. Assessment of Human Renal Transporter Based Drug-Drug Interactions Using Proximal Tubule Kidney-Chip, BioRxiv, 2022.

24. Arsenault P. et al. The Zinc Finger of Prolyl Hydroxylase Domain Protein 2 Is Essential for Efficient Hydroxylation of Hypoxia-Inducible Factor α, Molecular and Cellular Biology, 2016.

25. Jeewandara T. Experimental biology – Bioinspired engineering on a kidney chip to investigate renal calcification, Springer Nature Research Communities, Nature Portfolio, 2024.

26. Tang P. et al. Macrophages: versatile players in renal inflammation and fibrosis, Nature Reviews Nephrology, 2019.

27. Coralli C. et al. Alizarin Red Fluorescence Imaging for nano calcification, BioRxiv, 2022.

28. Khan S. et al. Kidney stones, Nature Review Disease Primers, 2016

29. Khan S. et al. Randall’s plaque and calcium oxalate stone formation: role for immunity and inflammation, Nature Reviews Nephrology, 2021.

30. Russel D. et al. The macrophage marches on its phagosome: dynamic assays of phagosome function, Nature Reviews Immunology, 2009

31. Vidarsson G. et al. IgG subclasses and allotypes: from structure to effector functions, Frontiers Immunology, 2014

32. Turner J. et al. Natural Killer Cells in Kidney Health and Disease, Frontiers Immunology, 2019

33. Duffield J. et al. Cellular and molecular mechanisms in kidney fibrosis, The Journal of Clinical Investigation, 2014

34. Zhou D. et al. Sonic hedgehog signaling in kidney fibrosis: a master communicator, Science China Life Sciences, 2017

35. Drobnik M. et al. Mechanosensitive Cation Channel Piezo1 Is Involved in Renal Fibrosis Induction, Molecular Science, MDPI 2024

36. Taguchi K. et al. Macrophage Function in Calcium Oxalate Kidney Stone Formation: A Systematic Review of Literature, Frontiers in Immunology, 2021

37. Khan S. et al. Association of Randall’s Plaques with Collagen Fibers and Membrane Vesicles, Journal of Urology, 2013

38. de Water R. et al. Calcium oxalate nephrolithiasis: effect of renal crystal deposition on the cellular composition of the renal interstitium, American Journal of Kidney Diseases, 1999

39. Murray P. et al. Macrophage activation and polarization: nomenclature and experimental guidelines, Immunity, 2014

40. Ingber D. E. et al. Tensegrity, cellular biophysics, and the mechanics of living systems, Reports on Progress in Physics, 2015

41. Negri A. Role of prolyl hydroxylase/HIF-1 signaling in vascular calcification, Clinical Kidney Journal, 2023

42. Ghafouri-Fard S. et al. A review on the role of cyclin dependent kinases in cancers, Cancer Cell International, 2022.

43. Sukov W. et al. CCND1 rearrangements and cyclin D1 overexpression in renal oncocytomas: frequency, clinicopathologic features, and utility in differentiation from chromophobe renal cell carcinoma, Human Pathology, 2009.

44. Hyndman K. Histone deacetylases in kidney physiology and acute kidney injury, Seminars in Nephrology, 2021.

45. Han W. et al. Hypoxia-inducible factor prolyl-hydroxylase-2 mediates transforming growth factor beta 1-induced epithelial–mesenchymal transition in renal tubular cells, Biochimica et Biophysica Acta (BBA) – Molecular Cell Research, 2013

46. Higgins D. et al. Hypoxia promotes fibrogenesis in vivo via HIF-1 stimulation of epithelial-to-mesenchymal transition, Journal of Clinical Investigation, 2007

47. Meneses A. et al. PHD2: from hypoxia regulation to disease progression, Hypoxia, 2016

48. Koul H. et al. Oxalate-induced initiation of DNA synthesis in LLC-PK1 cells, a line of renal epithelial cells, Biochemical and Biophysical Research Communications, 1994

49. Gaspar S. et al. Effect of calcium oxalate on renal cells as revealed by real-time measurement of extracellular oxidative burst, Biosensors and Bioelectronics, 2010

50. Thamilselvan S. et al. Free radical scavengers, catalase and superoxide dismutase provide protection from oxalate-associated injury to LLC-PK1 and MDCK cells, Journal of Urology, 2000

51. Koul H. et al. COM crystals activate the p38 mitogen-activated protein kinase signal transduction pathway in renal epithelial cells, Journal of Biological Chemistry, 2002

52. Hammes M. et al. Calcium oxalate monohydrate crystals stimulate gene expression in renal epithelial cells, Kidney International 1995

53. Thamilselvan S. and Menon M. Vitamin E therapy prevents hyperoxaluria-induced calcium oxalate crystal deposition in the kidney by improving renal tissue antioxidant status, BJU International, 2005

54. Coste B. et al. Piezo1 and Piezo2 are essential components of distinct mechanically activated cation channels, Science, 2010

55. He Y. et al. Myeloid Piezo1 Deletion Protects Renal Fibrosis by Restraining Macrophage Infiltration and Activation, Hypertension, 2022

56. Yang Y. et al. Functional Roles of p38 Mitogen-Activated Protein Kinase in Macrophage-Mediated Inflammatory Responses, Mediators of Inflammation, 2014

57. Raza A. et al. Anti-inflammatory roles of p38α MAPK in macrophages are context dependent and require IL-10, Journal of Leukocyte Biology, 2017

58. Cook-Mills J. et al. Vascular Cell Adhesion Molecule-1 Expression and Signaling During Disease: Regulation by Reactive Oxygen Species and Antioxidants, Antioxidants and Redox Signaling, 2011

59. Chiu J. et al. Shear stress increases ICAM-1 and decreases VCAM-1 and E-selectin expressions induced by tumor necrosis factor-[alpha] in endothelial cells, Atherosclerosis, Thrombosis and Vascular Biology (ATVB), 2003

60. Shinkai Y. and Tachibana M. H3K9 methyltransferase G9a and the related molecule GLP, Genes and Development, 2011

61. Yokoyama M. et al. Histone lysine methyltransferase G9a is a novel epigenetic target for the treatment of hepatocellular carcinoma, Oncotarget, 2017.

